# Distinct genetic origins of eumelanin intensity and barring patterns in cichlid fishes

**DOI:** 10.1101/2023.07.02.547430

**Authors:** A. Allyson Brandon, Cassia Michael, Aldo Carmona Baez, Emily C. Moore, Patrick J. Ciccotto, Natalie B. Roberts, Reade B. Roberts, Kara E. Powder

## Abstract

Pigment patterns are incredibly diverse across vertebrates and are shaped by multiple selective pressures from predator avoidance to mate choice. A common pattern across fishes, but for which we know little about the underlying mechanisms, is repeated melanic vertical bars. In order to understand genetic factors that modify the level or pattern of vertical barring, we generated a genetic cross of 322 F_2_ hybrids between two cichlid species with distinct barring patterns, *Aulonocara koningsi* and *Metriaclima mbenjii*. We identify 48 significant quantitative trait loci that underlie a series of seven phenotypes related to the relative pigmentation intensity, and four traits related to patterning of the vertical bars. We find that genomic regions that generate variation in the level of eumelanin produced are largely independent of those that control the spacing of vertical bars. Candidate genes within these intervals include novel genes and those newly-associated with vertical bars, which could affect melanophore survival, fate decisions, pigment biosynthesis, and pigment distribution. Together, this work provides insights into the regulation of pigment diversity, with direct implications for an animal’s fitness and the speciation process.

## INTRODUCTION

Coloration of animals has long fascinated both scientists and non-scientists alike. For centuries, scientists have asked questions about the underlying genetic control, diversity of variation, and ecological relevance of changes in pigmentation (Castle & Allen, 1903; Darwin, 1871; Wright, 1917). The rich collection of hues, spots, stripes, and bars of animals integrate both natural and sexual selective pressures (Brandon, Almeida, & Powder, 2023; Hoekstra, 2006; Hubbard, Uy, Hauber, Hoekstra, & Safran, 2010; Maan & Sefc, 2013). Pigmentation patterns are related to crypsis and predator avoidance, mate choice, color-mediated aggression, social dominance and competitive interactions, and collective animal behaviors such as schooling or shoaling in fishes (Brandon et al., 2023; Cuthill et al., 2017; Eizirik & Trindade, 2021; Hubbard et al., 2010; Korzan & Fernald, 2007; Maan & Sefc, 2013; Parichy, 2021; Protas & Patel, 2008; Sefc, Brown, & Clotfelter, 2014). Through this, these traits directly affect reproductive success, fitness, and speciation (Wagner, Harmon, & Seehausen, 2012), and the ultimate result is an incredible array of color pattern variation across animals.

One clade with notable variation in pigmentation is cichlid fishes, which have undergone a rapid and extensive adaptive radiation (Powder & Albertson, 2016; Santos, Lopes, & Kratochwil, 2023). Cichlids exhibit dramatic variation in their coloration, with variation due to species, sex, and geography (Konings, 2016; Maan & Sefc, 2013). The evolution of pigmentation is particularly important in cichlids, where sexual selection on divergent nuptial coloration appears to maintain pre-mating reproductive isolation among the most recently evolved species (Danley & Kocher, 2001). Much work has been done to begin to understand the molecular origins of the rich palette found across cichlids. Various genomic approaches have identified genetic loci that regulate black blotches (Roberts, Moore, & Kocher, 2017; Roberts, Ser, & Kocher, 2009), dark horizontal stripes (Kratochwil et al., 2018), yellow egg spots (Salzburger, Braasch, & Meyer, 2007; Santos et al., 2014), black and yellow coloration of the fins (Ahi & Sefc, 2017; O’Quin, Drilea, Conte, & Kocher, 2013), golden morphs (Wang, Xu, Kocher, Li, & Wang, 2022), albinism (Kratochwil, Urban, & Meyer, 2019), and even modularity in patterns across the flank (Albertson et al., 2014). However, one pigment phenotype that is understudied is the most common pigment pattern, dark vertical barring (Santos et al., 2023). In contrast to horizontal stripes, whose presence and absence is controlled by a master switch gene *agouti-related peptide 2* (*agrp2,* also known as *asip2b*) (Kratochwil et al., 2018), the presence of barring in cichlids is predicted to be polygenic (Gerwin, Urban, Meyer, & Kratochwil, 2021). These darkly pigmented bars are primarily due to a population of melanin producing cells called melanophores. Melanophores originate from trunk neural crest cells, as do other pigment cells in teleosts including xanthophores that generate red/yellow pigment and iridophores which are reflective (Parichy, 2021).

Here, we sought to determine the underlying genetic regulators of variation in vertical bar pigmentation. To accomplish this, we generated a genetic mapping cross of two Lake Malawi cichlids with alternate barring phenotypes. *Aulonocara koningsi* has high-contrast bars across its body, and *Metriaclima mbenjii* has fewer and fainter bars, with little contrast between bars and the background pigment levels (Figure 1a-b). By crossing two species that both display vertical barring, we set out to identify factors that alter the intensity and spacing of these bars, rather than master regulators governing their presence. In particular, we expected that one set of genomic regions would regulate where melanophores were located and the pattern of the bars, and a separate set of genes would independently regulate the levels of black/brown eumelanin being produced and dispersed from melanophores. We identify genetic intervals with candidate genes that are redeployed across vertebrates to regulate barring as well as other pigment phenotypes, as well as a series of additional genetic regions that are novel regulators of barring. Together, these data provide insights into the genetic and molecular underpinnings of pigment biodiversity, which lies at the intersection of a series of selective pressures that shape an animal’s ecology and evolution.

**Figure 1.**
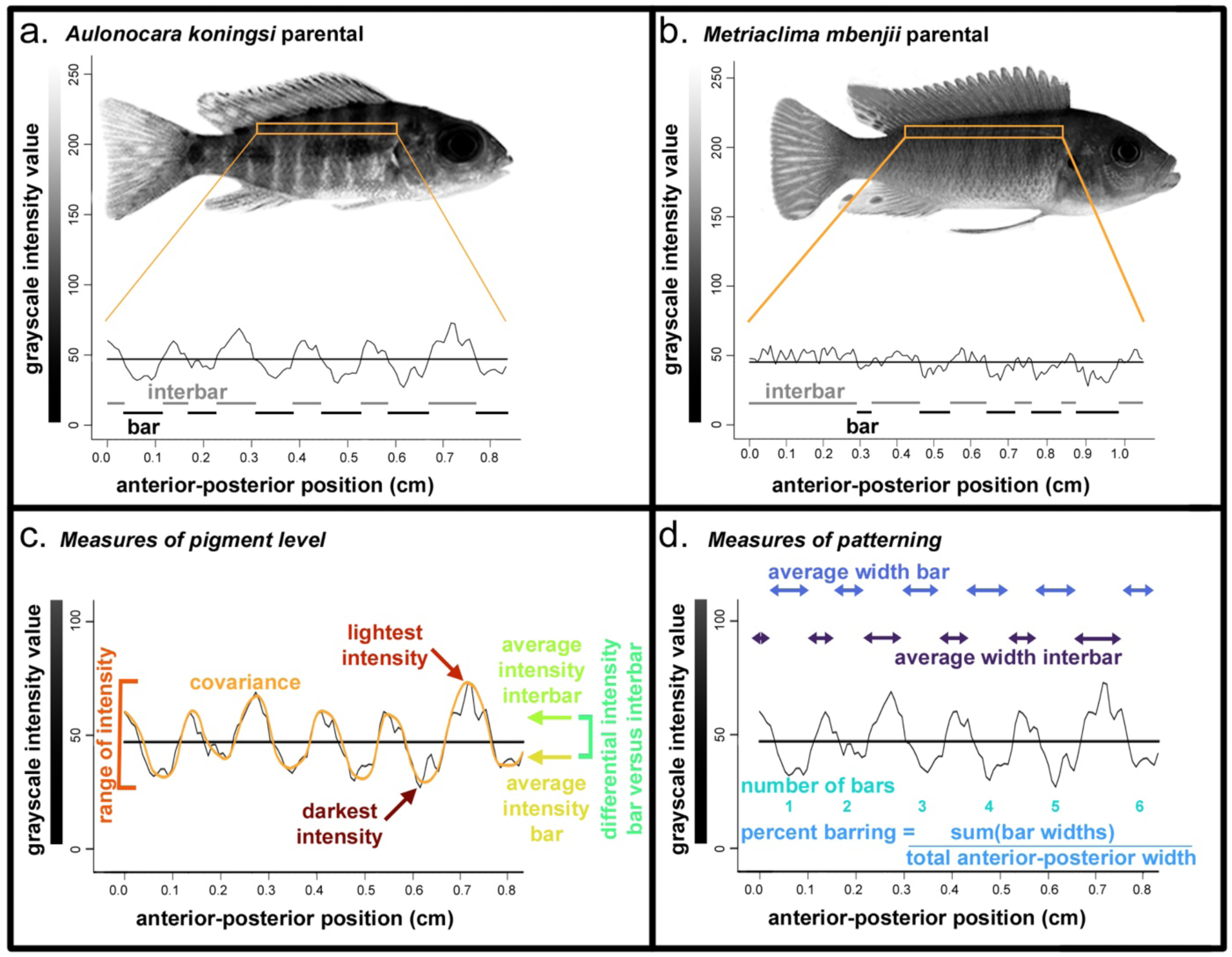
Parental species and measures of variation in barring pattern and pigment level. Representative (a) *Aulonocara koningsi* and (b) *Metriaclima mbenjii* parental, including quantification of region in orange rectangle into grayscale values. The horizontal bar in graphs in (a) and (b) are the average grayscale intensity value, which was calculated for each individual and used to characterize bar and interbars, indicated by black and gray marks, respectively, in (a) and (b). From grayscale plots for each individual, measures of (c) eumelanin pigment level and (d) bar patterning were calculated as visualized. Colors in (c) and (d) match colors used in Figure 3.

## MATERIALS AND METHODS

### Experimental cross

All animal care was conducted under approved IACUC protocol 14-101-O at North Carolina State University. A hybrid cross was generated from a single female *Metriaclima mbenjii* that was crossed to two male *Aulonocara koningsi*. The inadvertent inclusion of two grandsires resulted from an unexpected fertilization, as these animals externally fertilize. We discuss how this was accounted for during genotyping in the section *Genotyping and linkage map generation*. A single F_1_ family from this cross was subsequently in-crossed to generate an F_2_ hybrid mapping population. F_2_ hybrids were raised in density-controlled aquaria and with standardized measured feedings until around sexual maturity (five months of age), at which time they were sacrificed for analysis. The sex of each animal was determined based on a combination of gonad dissections at sacrifice and genotype at an XY locus on LG7 that solely determined sex in this cross (Peterson, Cline, Moore, Roberts, & Roberts, 2017; Ser, Roberts, & Kocher, 2010). Sex was omitted for animals with ambiguity or discrepancies between these calls (8.77% of animals), resulting in a set of 479 hybrids with 48.74% females.

### Imaging and filtering of data set

Images were taken of animals that were freshly sacrificed via cold buffered 100 mg/L MS-222. Euthanasia in cold solution relaxed chromatophores in the skin to maximize black/brown eumelanin-based pigmentation (Albertson et al., 2014). Whole fish photographs were taken using a uniform setup, with standard lighting conditions in a lightbox with a mirrorless digital camera (Olympus). All images included a scale bar and a gray scale color standard. Images were color balanced in Adobe Photoshop (version 22.0.0 or after) using the black and white segments of the color standard. From the total data set of 10 parentals of *Aulonocara koningsi*, 10 parentals of *Metriaclima mbenjii*, and 479 F_2_ hybrids, we omitted fish that exhibited two pigmentation phenotypes. First, *Metriaclima mbenjii* has a high percentage of animals that carry the ‘orange blotch’ (OB) phenotype, which results in marbled melanophore blotches rather than distinct barring (Konings, 2016; Roberts et al., 2017; Roberts et al., 2009). To enable analysis of barring patterns, we therefore removed 4 OB *Metriaclima mbenjii* parentals and 96 OB F_2_ hybrids. We further removed 65 hybrids that were heavily melanic to the degree that the eye was not distinguishable from the head or flank and thus anatomical landmarks used for additional processing were not visible. The final data set after this filtering included 10 *Aulonocara koningsi* parentals, 6 *Metriaclima mbenjii* parentals, and 322 F_2_ hybrids (48.97% female).

### Isolation and quantification of pigmented region

Images were rotated so a horizontal guideline aligned the midline of the caudal peduncle and the tip of the snout. A second horizontal guideline was added at the top of the caudal peduncle. From this image, a region was extracted for the remainder of the analysis (indicated by orange outlines in Figure 1a-b). This region was 10 pixels high with the ventral side aligned with the guide at the top of the caudal peduncle, the opercle at the anterior end, and the dorsal fin on the posterior end (Figure 1a-b). This standardized region of the body was chosen as it has a barring pattern representative of the entire flank and avoids areas that included the pectoral fin in a portion of images, which introduced variation in measurements of pigmentation.

The image of the isolated region was uploaded to FIJI software (version 2.9.0) (Schindelin et al., 2012), where it was converted to 32-bit grayscale. Following (Greenwood et al., 2011; O’Quin, Drilea, Roberts, & Kocher, 2012), the Plot Profile command in FIJI was used to convert the image to a numerical gray value from 0 (pure black) to 255 (pure white), averaging the values of the 10 pixels in each column (Figure 1). Isolated regions had an average width of 174 ± 36 pixels, which equated to 1.26 ± 0.26 centimeters or 30.6 ± 3.8 % of the total length (snout to caudal peduncle) of the animal.

### Quantification of melanic traits

Outputted data from Plot Profile in FIJI were analyzed in either R or with a custom perl script available at https://github.com/kpowder/Biology2022. The perl script used two criteria to define bars and interbars (defined as the region between bars) (Figure 1a-b) based on empirical testing of four *Aulonocara koningsi* parentals, four *Metriaclima mbenji* parentals, and four F_2_ hybrids, all randomly-chosen. Both cutoffs described below were selected as they accurately represented the barring pattern that was observed by eye on this test data set (Figure S1). First, we used the average intensity value to define bar regions, with gray intensity value less than (i.e., darker than) the average considered within a bar, and gray intensity value greater than this average considered within an interbar (Figure 1a-b and Figure S1). Second, to minimize overcounting of bars due to variation in pigment intensity from one pixel to the next, we required a bar to have at least 5 sequential pixels with intensity values below the average, and define the end of the bar as 5 pixels in a row above the average gray intensity value.

From the Plot Profile data and output of the perl script, we calculated seven measures related to variation in the levels of eumelanin produced: darkest intensity, lightest intensity, range of intensity, covariance of intensity measure with anterior-posterior position, the average intensity of bars, the average intensity of interbars, and the differential intensity between bars and interbars (Figure 1c). We note that an increased range of intensity, covariance, and differential intensity between bars and interbars are characteristic of animals with more discrepancy between bars and interbars (that is, highly melanic bars on very pale backgrounds). We also calculated four measures based around variation in the pattern of what regions of the body had bars: the total number of bars, the average width of bars, the average width of interbars, and the percent of barring, measured as the total length of regions classified as bars divided by the total length of the isolated region (Figure 1d).

### Statistical analysis of pigment measures

For each individual, the standard length of the animal was measured in FIJI as the number of pixels between the snout and the caudal peduncle, which was converted into absolute length in centimeters using a scale bar included in each picture. To remove the effects of allometry on pigment phenotypic measures, all measurements were converted into residual data by normalizing to the standard length, using a dataset including both parentals and hybrids. Analyses including linear normalization, ANOVAs, Tukey’s Honest Significant Difference post-hoc tests, and Pearson’s correlations were conducted in R (version 3.5.2 or higher).

### Genotyping and linkage map generation

Isolation, sequencing, and genotype calls are fully described in (DeLorenzo et al., 2023). Briefly, genomic DNA was extracted from caudal fin tissue, used to generate ddRADseq libraries, and sequenced on an Illumina HiSeq with 100bp paired end reads. Following demultiplexing and filtering of low-quality reads, reads were aligned to the *Metriaclima zebra* UMD2a reference genome and genotypes were called for those markers that had alternative alleles between the parents (i.e., AA x BB) and had a stack depth of 3. As mentioned above, an inadvertent fertilization event led to two grandsires in this cross. To focus on species-level genetic contributions, markers were excluded if *Aulonocara* sires had discrepant genotypes or Hardy-Weinberg equilibrium was not met. Any phenotypic effects of genetic variation from a single grandsire (that is, intraspecies variation) is expected to be diluted in this cross and therefore not be identified in the subsequent QTL mapping described in the following section.

Generation of the linkage map is fully described in (DeLorenzo et al., 2023), was built in R (version 4.0.3) with the package R/qtl (version 1.44-9) (Broman, 2009), and used custom scripts available at https://github.com/kpowder/Biology2022. Briefly, RAD markers were initially binned into linkage groups according to their position in the *M. zebra* UMD2a reference genome, cross referenced based on segregation patterns and recombination frequencies, and removed if located in unplaced scaffolds that had more than 40% of missing data or did not demonstrate linkage disequilibrium with multiple markers in a single linkage group. Additional manual curation was used to minimize the number of crossovers for those markers whose recombination frequency profile did not match their position in the linkage map, likely due to being within a misassembled region of the reference genome or a structural variant. The final genetic map included 22 linkage groups, 1267 total markers, 19-127 markers per linkage group, and was 1307.2 cM in total size.

### Quantitative trait loci (QTL) analysis

QTL analysis used the R/qtl package (version 1.44-9) (Arends, Prins, Jansen, & Broman, 2010; Broman, Wu, Sen, & Churchill, 2003) following (Jansen, 1994). Scripts are described in (Powder, 2020) and available at https://github.com/kpowder/MiMB_QTL. A multiple-QTL mapping (MQM) approach was used to more accurately identify intervals and their effects (Jansen, 1994).The approach starts by using the onescan function in R/qtl (Broman, 2009) to identify putative, unlinked QTL. These putative QTL were used as cofactors to build a statistical model, and were verified by backward elimination to generate the final model. Statistical significance was assessed using 1000 permutations to identify 5% (significant) and 10% (suggestive) cutoffs. For each of these QTL peaks, 95% confidence intervals on each linkage group were identified by Bayes analysis. Table S1 includes for each trait the cofactors used to build models, significance cutoffs, confidence intervals, and allelic effects at the peak marker of the QTL interval.

### Identification of candidate genes

Markers are named based on physical locations (contig and nucleotide position) in the *Metriaclima zebra* UMD2a reference genome. These nucleotide positions were used in the NCBI genome data viewer (https://www.ncbi.nlm.nih.gov/genome/gdv, *M. zebra* annotation release 104) to retrieve candidate Entrez gene IDs and genomic locations. If the extremes of the 95% confidence interval included markers that mapped to unplaced scaffolds, the closest marker that mapped to a placed scaffold was used as an alternative. Full gene names were extracted from the Database for Visualization and Integrated Discovery (DAVID) (Huang, Sherman, & Lempicki, 2009; Huang, Sherman, & Lempicki, 2009) using Entrez gene ID numbers as a query. Genes previously associated with body pigmentation or melanocyte development were extracted from the annotated Molecular Signatures Database (Liberzon et al., 2011), which is used for Gene Set Enrichment Analysis (Subramanian et al., 2005). A total of 258 genes from the human data set were used, which associated with the gene ontology (GO) cellular component term pigment granule (GO:0048770) and the following biological process terms: cellular pigmentation (GO:0033059), developmental pigmentation (GO:0048066), establishment of pigment granule localization (GO:0051905), melanocyte differentiation (GO:0030318), melanocyte proliferation (GO:0097325), melanosome assembly (GO:1903232), pigment accumulation (GO:0043476), pigment biosynthetic process (GO:0046148), pigment catabolic process (GO:0046149), pigment cell differentiation (GO:0050931), pigment granule localization (GO:0051875), pigment granule maturation (GO:0048757), pigment granule organization (GO:0048753), pigment metabolic process (GO:0042440), pigmentation (GO:0043473), positive regulation of developmental pigmentation (GO:0048087), positive regulation of melanocyte differentiation (GO:0045636), regulation of melanocyte differentiation (GO:0045634), regulation of pigment cell differentiation (GO:0050932), and regulation of pigmentation (GO:0120305).

## RESULTS

### Phenotypic variation in barring

We sought to examine the genetic factors that control variation in vertical bars across the flank, particularly the levels of eumelanin produced by melanophores and the spacing of bars and interbars. To accomplish this, we generated a hybrid cross between two species with distinct barring patterns. Importantly, given that both parents demonstrate some degree of barring, this cross is unlikely to identify genomic regions that are master regulators of bars (i.e., presence versus absence of bars), but rather how bars can be modified when they are present.

*Aulonocara koningsi* are distinguished by a regular series of vertical, melanic bars that extend on the anterior-posterior axis from the opercle to the caudal peduncle, and throughout the dorsal-ventral axis (Figure 1a). Melanic bars in *Aulonocara koningsi* are broken up by pale, lightly pigmented interbars that tend be narrower or the same width as melanic bars (Figure 1a). Alternatively, *Metriaclima mbenjii* have vertical bars that typically occur on the flank only between the opercle and anal fin, then become more irregular or stop towards the posterior of the animal (Figure 1b). Interbars on *Metriaclima mbenjii* are more melanic, such that the overall effect of barring in this species is a subtle vertical bar on a dark background (Figure 1b). In agreement with these qualitative observations, quantification of the level of melanic pigmentation reveals that compared to *Metriaclima mbenjii,* black or brown pigment in *Aulonocara koningsi* is not significantly darker (Figure 2a) but can be significantly lighter (Figure 2b). This generates a significantly larger range of pigment (Figure 2c) and larger differences between pigment levels in bars and interbars (Figure 2g).

**Figure 2.**
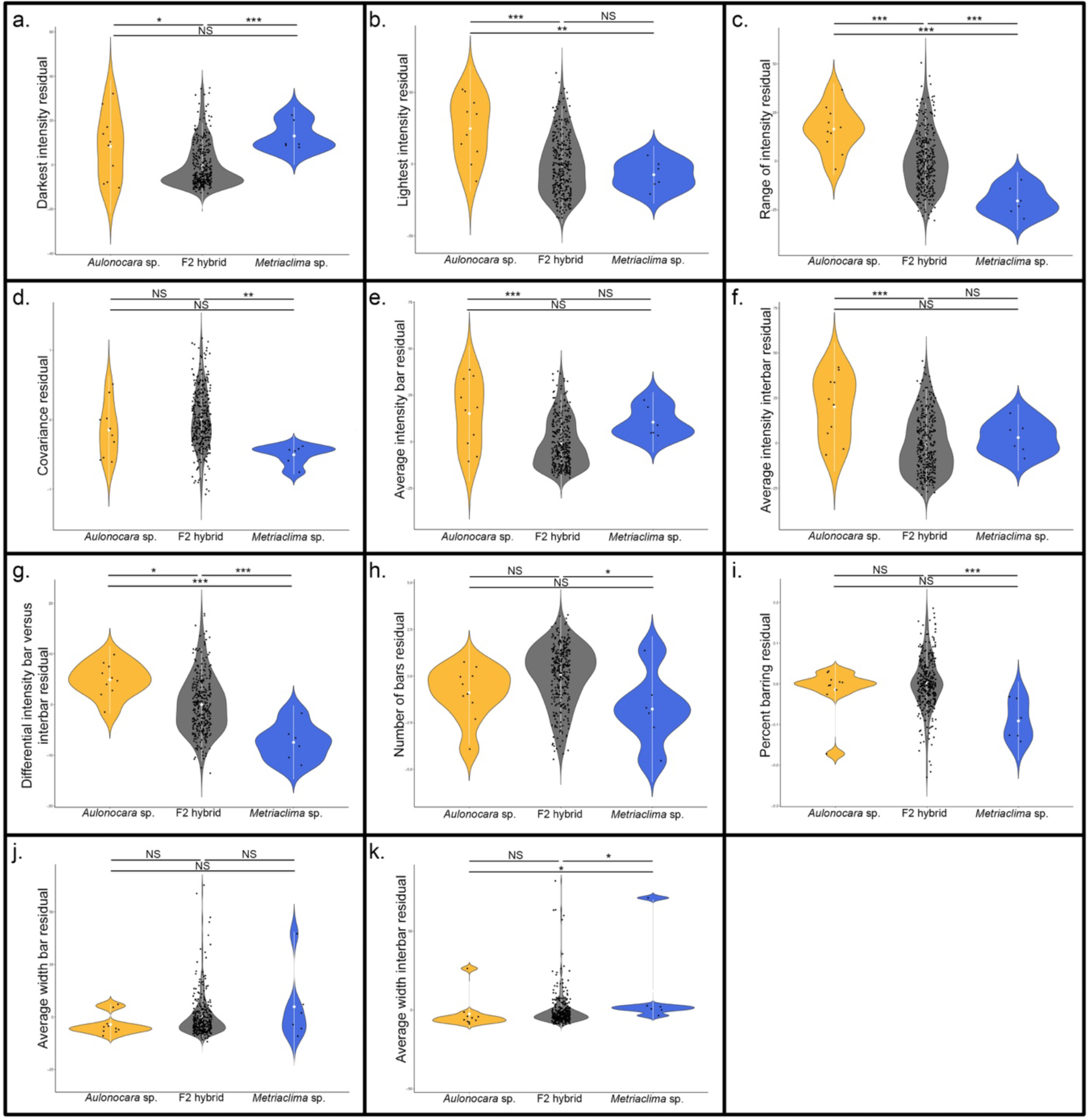
Variation in barring levels and patterns among *Aulonocara koningsi*, *Metriaclima mbenjii*, and their F_2_ hybrids. One set of measures relates to pigment levels produced by melanophores and are (a) darkest intensity, (b) lightest intensity, (c) range of intensity, (d) covariance, (e) average intensity of bars, (f) average intensity of interbars, and (g) differential intensity bars versus interbars. A second set of measures relates to the pattern of the bars and are (h) the number of bars, (i) percent barring, calculated as sum of total width of bars divided by total width of the isolated region, (j) average width of bars, and (k) average width of interbars. Significance in violin plots is based on ANOVA analysis followed by Tukeys HSD (data in Table S2; p-values indicated by * <0.05, ** <0.01, *** <0.005).

Though they have a visual difference in barring patterns, we did not find that parental species were significantly different in their patterns of vertical bars (Figure 2h-j). The exception to this is that *Metriaclima mbenjii* had significantly wider interbars (Figure 2k), though this is likely driven by the fact that the posterior of the flank in this species often did not have distinct bars, and due to a single specimen that had only a single bar and nearly the whole length of the body was classified as an interbar. We expect two factors explain the lack of statistical significance for most of the traits examined between the parental species. First, *Metriaclima mbenjii* is noted for a high percentage of ‘orange blotch’ (OB) animals, where melanophores are organized in irregular patches rather than bars. After removing these animals from the data set in order to focus on barring phenotypes, this only left six *Metriaclima mbenjii* parental specimens, reducing our statistical power. Second, our analysis that classified bars and interbars can have errors in classification for an animal in which the grayscale intensity has less range and more inconsistent fluctuations, which was observed in many of the *Metriaclima mbenjii* parentals (i.e., compare pattern of the graph in Figure 1a versus variation around the average value in Figure 1b). An inability to accurately define bars and interbars in *Metriaclima mbenjii* parentals would influence measures of the average intensity of bars, the average intensity of interbars, the differential intensity between bars and interbars, the number of bars, the percent of barring, the average width of bars, and the average width of interbars, most of which did not show significant differences between parentals (Figure 2e-k).

Despite this, it’s important to note that 100% of the 322 F_2_ hybrids demonstrated a distinct barring pattern, resembling the *Aulonocara koningsi* parental phenotype, and thus would be classified correctly by our analysis approach. The dominance of this overall barring pattern observed in the *Aulonocara* x *Metriaclima* F_2_ hybrids agrees with a previous suggestion that several genes are sufficient to drive the formation of bars (Gerwin et al., 2021). Though they all had melanic vertical bars, F_2_ hybrids were phenotypically varied in terms of the level of eumelanin produced in the bars and interbars, as well as the spacing of these pigment elements. This population therefore can yield valuable insights into the patterns of phenotypic variation in barring, as well as the genetic loci that regulate this.

Color differences between males and females is thought to be sexually-selected, resulting in widespread sexual dimorphism across vertebrates (Bell & Zamudio, 2012; Hubbard et al., 2010; Miller, Mesnick, & Wiens, 2021; Williams & Carroll, 2009), including cichlids (Brzozowski, Roscoe, Parsons, & Albertson, 2012; Konings, 2016; Salzburger, 2009; Santos et al., 2023). Within this cross however, we find no statistical relationship between any measures of pigment level or patterns and sex (p = 0.24 to 0.89, Table S2), nothing that F_2_ animals were collected as juveniles and did not express fully mature nuptial coloration. Thus, variation identified here reflects differences due to species-specific genetic polymorphisms. Notably, within Lake Malawi cichlids, a set of ancestral polymorphisms are being recombined in differing combinations among species (Brawand et al., 2014; Malinsky et al., 2018; Svardal et al., 2020). Thus, even for traits with non-significant differences between parental species, QTL mapping can identify genetic factors that underlie pigment variation within this radiation and genetic combinations that are possible in other species.

### Genetic basis of variation in pigment phenotypes

To determine the genetic basis of variation in barring phenotypes, we genetically mapped seven traits related to the level of eumelanin produced and four traits related to the location of bars. Fifty-one QTL underlie quantitative differences in pigmentation in *Metriaclima* x *Aulonocara* F_2_ hybrids, with 48 that reach 5% statistical significance at the genome-wide level (Figure 3, Figure S2, Figure S3, and Table S1). An additional 3 QTL are suggestive at the 10% level and included in supplemental material only (Figure S2, Figure S3, and Table S1). QTL are found on 17 of 22 linkage groups, with each linkage group containing 1-6 significant loci each.

**Figure 3.**
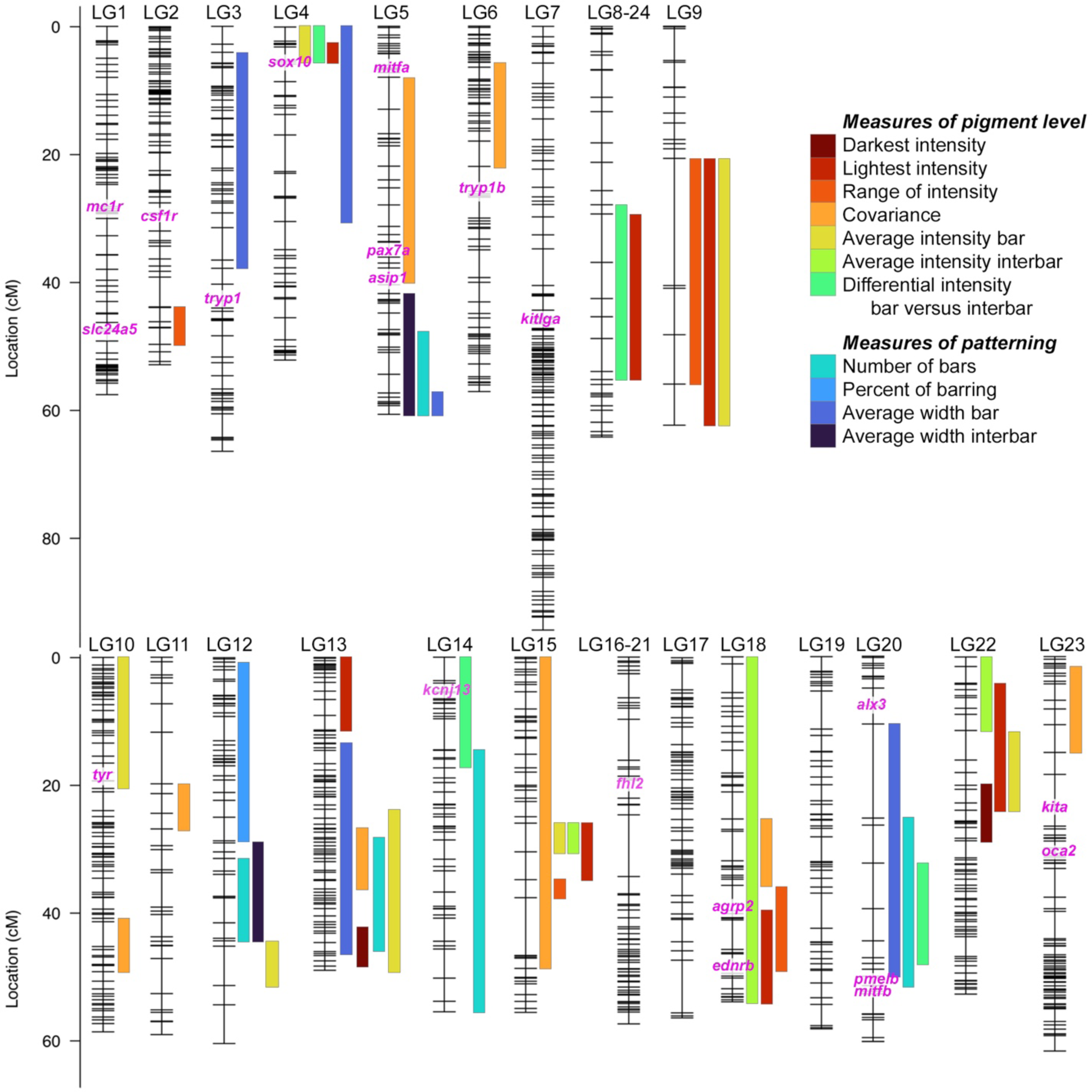
Quantitative trait loci (QTL) mapping identifies 48 intervals associated with variation in barring between *Metriaclima mbenjii* and *Aulonocara koningsi*. Each linkage group (LG, i.e., chromosome) has markers indicated by hash marks. Bar widths indicate 95% confidence interval for each QTL and bar color indicates the pigment trait analyzed. Candidate genes previously associated with variation in eumelanin production and development of stripes or bars (see main text for references) are in pink text, with their genomic locations indicated on linkage groups. Additional candidate genes *pax3a* and *pmela* are located in unplaced scaffolds in the *M. zebra* UMD2a reference genome and not included here. Illustrations of each trait are in Figure 1. QTL scans at the genome and linkage group level are in Figures S2 and S3, respectively. Details of the QTL scan, including statistical model and physical locations defining each QTL are in Table S1.

Between one and six QTL contribute to each trait, with each interval explaining 4.75-16.1% of variation for each trait (average 7.25% variation explained, Figure S3 and Table S1). While these data indicate that all these pigment traits are multifactorial, the QTL identified cumulatively explain a considerable portion of variation for multiple traits. Five QTL together explain 42.66% of variation in the number of bars, eight QTL explain 44.07% of variation in the average intensity of bars, seven QTL combine to explain 50.61% of variation in lightest intensity, and eight QTL cumulatively explain 69.14% of the total variation in covariance, or the discrepancy between dark bars on a lightly pigmented flank.

There is a large degree of overlap in our QTL intervals, which is expected given both the traits analyzed and their degree of correlation. For example, LG9 contains three QTL that overlap between 20.7-56.0 cM and control the lightness intensity, range of intensity, and average intensity of bars (Figure 3 and Table S1). A change in lightness would directly affect the calculation of the range of intensity, explaining why these traits map to the same interval. Accordingly, these traits are highly correlated (r = 0.90, Table S3) and the lightest intensity is also correlated with the average intensity of bars (r = 0.87, Table S3). These types of phenotypic correlations also explain the overlapping QTL on LG15 from 26.0-30.5 cM for lightest intensity, average intensity of bars, and average intensity of interbars (Figure 3 and Table S1), traits that all correlate (r = 0.87 to 0.96, Table S3). However, it’s important to note that even highly correlated traits can be regulated through distinct mechanisms (i.e., many-to-one mapping (Wainwright, Alfaro, Bolnick, & Hulsey, 2005)). For example, darkest intensity, average intensity of bars, and average intensity of interbars are all highly correlated (0.84 < r < 0.95, Table S3). Together, these traits map to fourteen intervals, ten of which are unique genomic regions. Further, five of the eight QTL for the average intensity of bars are on LGs that do not contain any QTL for the other two correlated traits.

The effect of specific alleles on phenotypes is varied across phenotypes (Figure S3 and Table S1). For instance, for the QTL on LG12, the allele inherited from the *Metriaclima mbenjii* granddam decreases the number of bars on LG12, but increases the number of bars on LG13. Three other examples highlight the complex genetic interactions that are possible within these species and across the cichlid radiation. First are the set of overlapping QTL on LG9 and cluster of QTL on LG22. In both cases, alleles inherited from *Metriaclima mbenjii* are associated with lighter intensity values (Figure S3 and Table S1), though lighter pigmentation is associated with the *Aulonocara koningsi* phenotype (Figure 1-b and Figure 2b). The second example is a group of QTL related to pigmentation levels that occur on LG15. All 5 QTL demonstrate an underdominant inheritance pattern, such that a combination of heterozygous alleles explain the variation in lightest intensity, range of intensity, covariance, average intensity of bars, and average intensity of interbars (Figure S3 and Table S1). These suggest that complex interactions between genes (i.e., epistasis (Carlborg & Haley, 2004; Phillips, 2008)) regulate these phenotypic traits such that the effects of the *Metriaclima mbenjii* allele are only visible in the absence of or in combination with additional alleles. These examples of cryptic genetic variation (Gibson & Dworkin, 2004; Paaby & Rockman, 2014) are important to understand the full spectrum of genetic factors that regulate a complex trait like pigmentation, and allelic combinations that are likely to be present in other cichlid species within Lake Malawi (Brawand et al., 2014; Malinsky et al., 2018; Svardal et al., 2020).

### Distinct genetic origins underlie pigment levels and patterning

We hypothesized that a separate set of genetic, molecular, and developmental mechanisms may underlie variation in the level of melanic pigmentation and the patterns of vertical bars. That is, we predicted that one set of genes would control where melanophores would be located and capable of generating dark vertical bars (i.e., patterning), and a separate set of genes would then control how much eumelanin is produced by these melanophores (i.e., pigment level). One set of data supporting this is an analysis of correlations among traits (Table S3). While some measures of pigment levels are correlated as discussed above, none of the four measures related to patterning were correlated with any of the seven measures of pigment level (r = −0.39 to 0.31 in pairwise comparisons, Table S3).

QTL data also largely supports that divergent genetic factors regulate pigment intensity and bar/interbar locations, through the examination of the degree of overlap of QTL intervals (Figure 3). We found numerous overlaps within similar types of traits. That is, LG5 and LG12 contain regions that have 2-3 overlapping QTL related to patterning, and LG8-24, LG9, LG15, LG18, and LG22 have 2-4 QTL at the same genomic locus that control levels of pigment produced.

Only LG4, LG13, LG14, and LG20 have overlapping 95% confidence intervals for traits related to pigment level and patterning, and additional examination of these regions suggest that a pleiotropic effect on both aspects of pigmentation is limited to LG13 and LG20 (Figure 4). For instance, all four QTL on LG4 have 95% confidence intervals that overlaps from 0-5.79 cM (Figure 3 and Table S1). However, this includes the peak and full region for the three traits related to pigment level, while the QTL for patterning has a peak at 20 cM and dips below significance under the peak of the other QTL, suggesting that two linked, but distinct loci may underlie variation in the two traits (Figure 4). Another overlap occurs on LG14, where a QTL for the differential intensity between bars and interbars overlaps with a QTL for the number of bars from 14.6-17.32 cM (Figure 3 and Table S1). However, closer examination of these loci reveals that the peak for these QTL are on opposite ends of the linkage group, 55 cM apart (Figure 4), and this overlapping interval is unlikely to contain a causative gene that underlies both the pigment level phenotype and the patterning phenotype.

**Figure 4.**
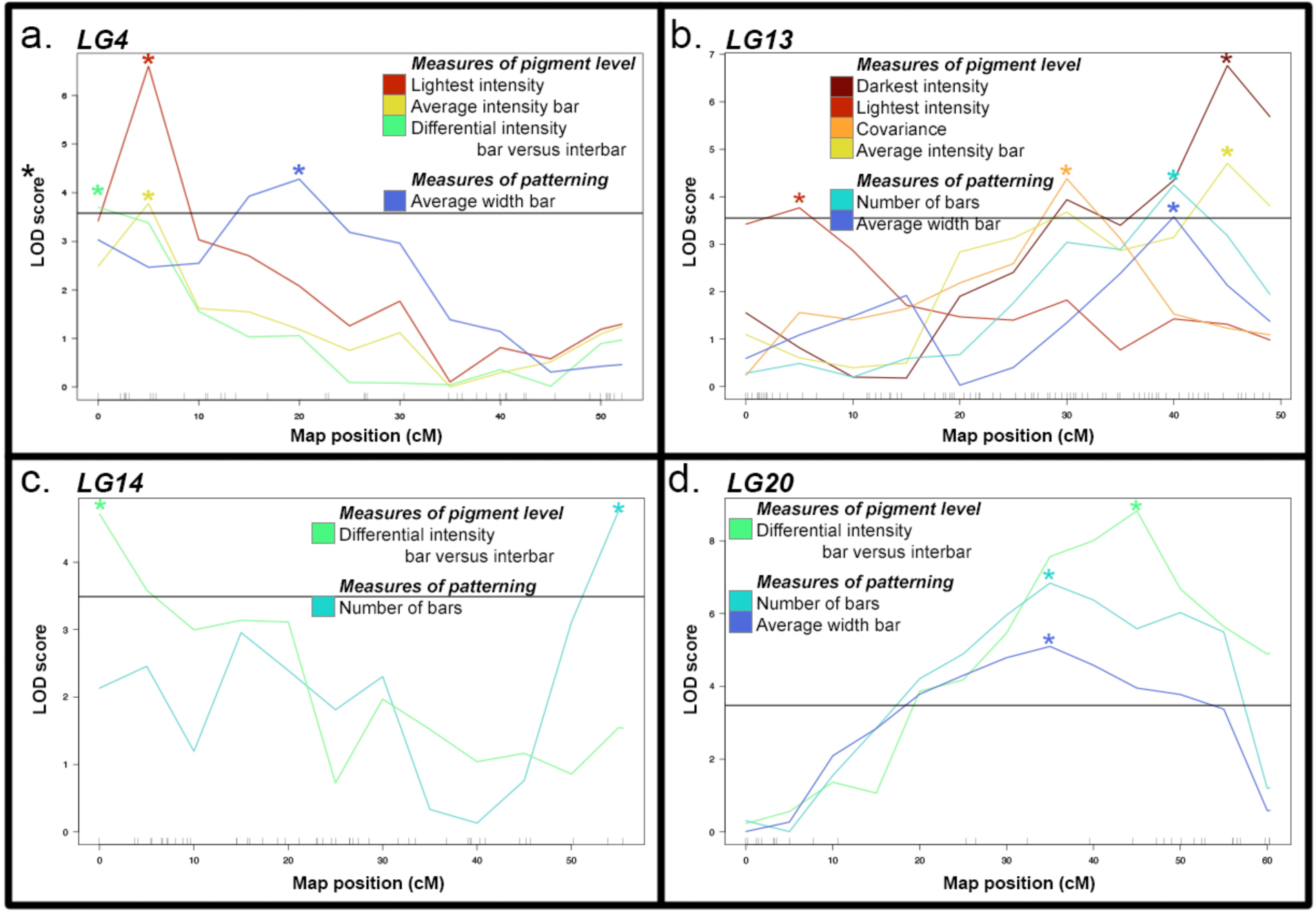
Quantitative trait loci (QTL) that underlie variation in pigment levels and pigment patterning are largely distinct. Included are all linkage groups—(a) LG4, (b) LG13, (c) LG14, and (d) LG20—in which QTL for pigment level and patterning have overlapping 95% confidence intervals as visualized in Figure 3. Colors represent trait, as indicated by the legend and as illustrated in Figure 1. Peak markers for each QTL are indicated by an asterisk in a color matching the trait. The solid horizontal line in each panel represents 5% significance, measured as the average value from each of the featured scans on that specific linkage group; averaging this significance did not cause any of these QTL to change from significant to non-significant or vice versa. Further details of the QTL are in Figure S3 and Table S1.

However, two regions may regulate both pigment intensity and location on the body. First is on LG20, where a QTL for differential intensity between bars and interbars resides within QTL for number of bars and average width of bars (Figure 3). While the peak for the patterning QTL are both at 35 cM and the peak for the pigment level QTL is at 45 cM, these three all feature broad peaks with a high degree of overlap (Figure 4 and Table S1). Finally, six separate QTL reside on LG13 and span the entirety of the chromosome (Figure 3). Examination of the peaks and confidence intervals (Figures 3-4 and Table S1) suggests that at least 3 separate regions of LG13 are contributing to the pigment traits measured. This includes a region from 0-11.37 cM containing a QTL for lightest intensity, a region from 26.11-36.24 cM with a QTL for covariance, and a region from 42.85-48.33 cM that includes a QTL for darkest intensity (Figures 3-4). These three QTL do not overlap each other, but the latter two both overlap QTL for average intensity of bars, number of bars, and average width of bar (Figures 3-4). Thus, in the case of both LG13 and LG20, more detailed mapping will be necessary to determine if these traits are regulated by the same gene, distinct genes that are in close physical distance or linkage with each other, or distinct genomic loci.

## DISCUSSION

Though it is one of the most common pigment patterns in fishes, we know relatively little about the genetic basis of dark vertical barring (Santos et al., 2023). To address this, we used a genetic mapping cross between two cichlids with distinct presentations of barring and mapped 48 genetic loci that influence the relative levels of melanic pigment in bars and interbars, as well as their patterning. In addition to identifying this series of quantitative trait loci, we found eumelanin pigment levels and barring patterns are largely regulated by independent loci. These separate, polygenic genetic architectures would enable evolutionary fine-tuning of barring in response to natural or sexual selection, promoting further diversity in pigmentation. Further, we observed that vertical barring is not due to a master regulatory gene, but a combination of genetic factors. This directly contrasts with two other traits with distinct arrangements of melanophores, horizontal stripes and blotching, controlled by *agouti-related protein 2* (*agrp2*) and *paired box 7a* (*pax7a*), respectively (Kratochwil et al., 2018; Roberts et al., 2009). Our data thus supports a previous suggestion that barring in cichlids is polygenic (Gerwin et al., 2021).

Our loci add to a variety of genetic factors of both small and large effect that regulate pigmentation in cichlids (Albertson et al., 2014; Kratochwil et al., 2018). These alleles for fin pigmentation (Ahi & Sefc, 2017; Albertson et al., 2014; Salzburger et al., 2007; Santos et al., 2014), xanthophore-based red and yellow coloration (Albertson et al., 2014; Wang et al., 2022), melanophore-based black and brown coloration (Albertson et al., 2014; Kratochwil et al., 2018; Roberts et al., 2009), and integration versus modularity of color patterns across the flank (Albertson et al., 2014) are shuffled in differing combinations (Brawand et al., 2014; Malinsky et al., 2018; Svardal et al., 2020) to generate the range of colors and patterns that characterize the adaptive radiation of cichlids (Kocher, 2004; Konings, 2016; Santos et al., 2023). These pigmentation patterns, whether inherited independently or not, are then subject to a variety of ecologically-relevant selective pressures such as predator avoidance and intrasexual competitive interactions (Brandon et al., 2023; Cuthill et al., 2017; Eizirik & Trindade, 2021; Hubbard et al., 2010; Korzan & Fernald, 2007; Maan & Sefc, 2013; Parichy, 2021; Protas & Patel, 2008; Sefc et al., 2014). One critical implication of these hues and patterns is assortative mating (Couldrige & Alexander, 2002; Jordan, Kellogg, Juanes, & Stauffer, 2003), which can directly result in reproductive isolation, and thus sexual selection has been central to the dramatic speciation and divergence of cichlids (Danley & Kocher, 2001; Muschick et al., 2014; Ronco et al., 2021; Wagner et al., 2012).

The majority of our QTL do not include a series of genes previously associated with variation in pigment (Figure 3) in cichlids (Albertson et al., 2014; Kratochwil et al., 2018; Kratochwil et al., 2019; Roberts et al., 2009; Salzburger et al., 2007; Santos et al., 2014; Wang et al., 2022), *Danio* species including those with vertical barring (Lamason et al., 2005; Mills, Nuckels, & Parichy, 2007; Parichy et al., 2000; Parichy, Rawls, Pratt, Whitfield, & Johnson, 1999; Parichy & Turner, 2003; Podobnik et al., 2020), cavefish (Gross, Borowsky, & Tabin, 2009; Protas et al., 2006), sticklebacks (Greenwood, Cech, & Peichel, 2012), and other non-fish vertebrates (Domyan et al., 2014; Hoekstra, 2006; Jablonski, 2021; Lamason et al., 2005; Lu et al., 2016; Mallarino et al., 2016). For instance, *melanocortin 1 receptor* (*mc1r*) on LG1 has been associated with a series of adaptive pigment changes, through regulation of the biosynthesis of eumelanin (Hoekstra, 2006). While activating or repressing eumelanin production would likely influence any of the seven traits related to pigment level that we measured, none of the 35 QTL that underlie these traits include *mc1r*. Additional work will be needed to narrow genetic intervals, verify candidate genes, and identify the molecular and cellular mechanisms that generate the phenotypes we mapped. Most of our genetic intervals contain many genes, from 40 genes for covariance on LG10 to the entire chromosome and 1069 genes for the average intensity of interbars on LG18 (average = 342 genes, Table S4). However, we discuss below a number of strong candidate genes within these intervals and how they may mediate variation in melanic traits.

One set of candidate genes are associated with the development and survival of the melanophores themselves, including trunk neural crest cells which are the embryonic source of these pigment cells (Brandon et al., 2023; Parichy, 2021). Variation in the induction, migration, and differentiation process could change the number and/or location of melanophores present within the skin to generate changes in both the pattern of barring and the intensity of melanin produced. For instance, a QTL for average width of bars on LG4 includes the candidate gene *SRY-box 10* (*sox10*). We note that *sox10* is near, but not included in QTL for the lightest intensity, average intensity of bars, and differential intensity between bars and interbars that partially overlap this QTL for average width of bars (Figure 3 and Table S1). *Sox10* is necessary for neural crest cell specification (Carney et al., 2006; Jacob, 2015) and required to establish the melanophore linage (Marathe et al., 2017). Genetic variation in this gene in humans results in Waardenburg syndrome, which is characterized by a suite of alterations to neural crest cell derivatives, one of which is depigmented patches in the skin and hair (Pingault et al., 2010; Pingault, Zerad, Bertani-Torres, & Bondurand, 2022). This is not the only candidate gene associated with a pigmentation condition in humans. Overlapping QTL on LG13 contribute to covariance, the number of bars, and the average width of bars (Figure 3). These three QTL all include the candidate gene *F-box protein 11a* (*fbxo11a*) and the QTL for number of bars and average width of bars also include *SPARC-related modular calcium binding 2* (*smoc2*) (Table S4). Both genes are associated with the human condition vitiligo, characterized by the progressive loss of pigment cells through cell death or autoimmunity (Alkhateeb, Al-Dain Marzouka, & Qarqaz, 2010; Birlea, Gowan, Fain, & Spritz, 2010; Le Poole et al., 2001; Xie et al., 2016). Though little is known about the molecular function of *smoc2*, *fbxo11* regulates melanocyte proliferation, apoptosis, and intracellular transport of the eumelanin biosynthesis enzyme tyrosinase (Guan et al., 2010). Such a loss of melanophores in localized regions could result in the changes in barring pattern or amount of melanin produced and counting of individual melanophores (O’Quin et al., 2013; O’Quin et al., 2012) may provide further insights into this regulation.

The location and number of melanocytes would also be impacted by genes that regulate fate decisions during melanophore development. The QTL for covariance on LG5 includes the gene *pax7a*, and a related gene, *paired box 3b* (*pax3b*), is located within a QTL on LG14 for differential intensity of bars and interbars. *Pax3a*, the paralog of our candidate, and *pax7a* and can act transcriptionally as switch factors between different pigment cell fates, and these genes have previously been associated in cichlids with changes in the balance of melanophore and xanthophore cell numbers, changes in pigment levels, or altered patterns such as melanic blotches (Albertson et al., 2014; Minchin & Hughes, 2008; Roberts et al., 2017; Roberts et al., 2009). An overlapping QTL for the lightest intensity and the average intensity of interbars on LG18 includes another candidate gene related to cell fate decisions. *Endothelin receptor type B* (*ednrb*) is required for differentiation of melanophores (Saldana-Caboverde & Kos, 2010) and another pigment cell in fishes, iridophores (Krauss et al., 2014). Mutations in *ednrb* result in broken stripes in zebrafish (Parichy et al., 2000), though it has yet to be determined if the spots caused by this fate switch could merge into a bar pattern instead of stripes. Also within this interval on LG18 is the master switch for horizontal stripes in cichlids, *agrp2* (Figure 3). Previous work has predicted that stripes and bars are regulated by genetically-independent modules (Gerwin et al., 2021), suggesting that *agrp2* is not the causative gene on LG18 for changes in our bar phenotypes. However, it is possible that this independence depends on the cichlid species being compared, and *agrp2* may regulate barring in *Metriaclima* and *Aulonocara*.

Once melanophores are specified and migrate to their position on the flank, variation in eumelanin biosynthesis can produce variation. This would be expected to change pigment intensity, but not the pattern of barring. In agreement with this, the three QTL intervals described below that contain candidate genes associated with eumelanin production are associated with at least one of the measures of pigment level, but none of the measures of bar location. The LG5 QTL for covariance and the LG10 QTL for average intensity of bars contain two genes that have been associated with the biosynthesis of melanin across multiple vertebrate species, *agouti signaling protein* (*asip*) and *tyrosinase* (*tyr*) (Hoekstra, 2006). Interestingly, *asip* is also associated with countershading, a dorsoventral gradient of pigmentation important for predator avoidance, the switch between production of dark eumelanin and yellow/red pheomelanin, and may also repress melanophore differentiation (Cal et al., 2019; Ceinos, Guillot, Kelsh, Cerda-Reverter, & Rotllant, 2015; Steiner, Rompler, Boettger, Schoneberg, & Hoekstra, 2009). Candidate genes may also regulate melanin production based on ecological triggers. For instance, within the LG15 QTL associated with lightest intensity, covariance, average intensity of bars, and average intensity of interbars is the candidate gene *melanocortin 2 receptor accessory protein 2* (*mrpa2*). This gene is involved in the melanocortin response pathway, and can modulate melanin levels following starvation- or crowding-induced stress (Cortes et al., 2014).

Finally, variation in barring can be generated by differences in pigment density and distribution or alteration of melanophore cell density and shape (Liang, Gerwin, Meyer, & Kratochwil, 2020). While this can be rapidly regulated in cichlids through physiological changes such as hormone signaling (Muske & Fernald, 1987; O’Quin et al., 2012), this can also be regulated at the genetic level. Within the overlapping regions on LG4 for QTL regulating lightest intensity, average intensity of bars, and differential intensity between bars and interbars is *melanin-concentrating hormone receptor 1b* (*mchr1b*). This G-protein coupled receptor integrates with the nervous system to regulate hormonal changes in pigment aggregation as a fish alters its pigmentation for camouflage from predators, to attract a mate, or in response to intrasexual competition (Madelaine, Ngo, Skariah, & Mourrain, 2020; Mizusawa et al., 2011). Another strong candidate gene is found within a QTL on LG14 for differential intensity between bars and interbars. *Potassium inwardly rectifying channel subfamily J member 13* (*kcnj13*) is necessary for interactions between melanophores and other pigment cell types such as iridophores and xanthophores, resulting in localized changes in chromatophore shape and changes in the contrast of pigment patterns (Podobnik et al., 2022). Notably, *kcnj13* is associated with the evolution of vertical barring in *Danio* species, suggesting a conserved role in barring across a large portion of the fish phylogeny (Podobnik et al., 2020). Finally, a QTL for the number of bars on LG20 includes the gene *premelanosome protein b* (*pmelb*). *Pmelb* encodes a protein specific to pigment cells, that affects the cellular structure and shape of melanosomes through formation of fibrillar sheets on which melanin polymerizes and is deposited (Hellstrom et al., 2011; Schonthaler et al., 2005; Watt, van Niel, Raposo, & Marks, 2013). Further, CRISPR inactivation of paralogs *pmela* and *pmelb* in tilapia resulted in a reduction of melanophore number and size, as well as a loss of a vertical barring pattern (Wang et al., 2022).

## CONCLUSIONS

Pigment hues and patterns can be selected by a series of natural and sexual selective pressures including predator avoidance, mate choice, and competitive interactions. Here we explore one common pattern with the diverse coloration found in cichlid fishes, vertical melanic barring, for which the genetic and molecular basis is largely unexplored. We show here that the genomic intervals that influence pigment levels are largely distinct from those that regulate bar patterning, which can promote the degree of variation that is possible in this trait. A series of candidate genes within these intervals highlight the varied ways that melanophore development can be altered to produce ecologically-relevant variation in barring. The pigmentation patterns studied here are particularly important for the adaptive radiation of cichlids, where they play a role in sexual selection and reproductive isolation, and therefore in maintaining species boundaries. Future studies identifying the causative alleles for the QTL we identify here will allow exploration of their evolutionary history across the cichlid radiation, and their potential role in speciation.

## ACKNOWLEDGEMENTS

This work was supported by NSF CAREER IOS-1942178 (KEP), NIH P20GM121342 (KEP), NSF IOS-1456765 (RBR), and an Arnold and Mabel Beckman Institute Young Investigator Award (RBR). This work is dedicated to the memory of Dr. Stephen L. Johnson, who first introduced KEP to the beauty found in fish melanophores, and whose work helped inspire RBR to enter the world of fish genetics.

## DATA ACCESSIBILITY

Raw sequence data are available at https://www.ncbi.nlm.nih.gov/bioproject/PRJNA955776. Additional data are available at Dryad [link to be provided prior to publication]. Dryad files include phenotypic measures and genotypes used for quantitative trait loci mapping.

## AUTHOR CONTRIBUTIONS

KEP and RBR designed the research. ACB, ECM, PJC, and NBR performed animal husbandry, photography, and collections. NBR prepared sequencing libraries. AAB, CM, ECM, ACB, RBR, and KEP analyzed data. AAB and KEP wrote the paper with edits from all authors.

## Notes

### Competing Interest Statement

The authors have declared no competing interest.

