## Supporting information for "Distinct genetic origins of eumelanin intensity and barring patterns in cichlid fishes"

Supplemental table legends and figures with legends for Brandon et al.

1. Legends for Supplemental tables, provided as separate .xlsx files (pg. 1)
    a. Table S1: Details of quantitative trait loci (QTL) mapping and regions
    b. Table S2: Statistical significance of pigment measures
    c. Table S3: Correlations between pigment traits
    d. Table S4: Candidate genes for pigment QTL
2. Figure S1: Determination of thresholds used to define bars (pg. 2)
3. Figure S2: Genome-wide QTL scans for pigment traits (pg. 3-4)
4. Figure S3: QTL scans by linkage group and allelic effects (pg. 5-22)

**Table S1. Details of quantitative trait loci (QTL) mapping and regions.** For each QTL, we include cofactors use to generate models as well as markers and physical positions for the peak of the QTL and the 95% confidence interval. Marker names include the physical location on the linkage group, with names referring to the contig and nucleotide position in the *M. zebra* UMD2a assembly. LOD values, percent phenotypic variance explained by that QTL, allelic effects, and additive, dominance, and heritability calculations are for the peak marker in the QTL. QTL listed in gray are suggestive at the 10% significance level, while those in black meet 5% genome-wide significance based on values indicated.

**Table S2. Statistical significance of pigment measures.** Effects of size (standard length) and sex were assessed by ANOVA analysis. Note that sex was only called for $F_2$ hybrids, but not parental species. Significance between parental and hybrid groups on size-corrected (i.e., residual values) were assessed by ANOVA followed by Tukeys HSD. P-values < 0.05 are in bold red text.

**Table S3. Correlations between pigment traits**. Calculations are for $F_2$ hybrids animals only and exclude parentals. Except for correlations with standard length, all measures are size corrected prior to the calculation of correlation. Values that are >0.80 or <-0.80 are in bold red text.

**Table S4. Candidate genes for pigment QTL.** For the indicated 95% confidence interval, we include candidate gene names in the *M. zebra* annotation release 104, NCBI gene ID number, and physical positions in the *M. zebra* genome UMD2a assembly. Those genes that have been previously associated with pigmentation or melanocyte development or function in the Molecular Signatures Database/Gene Set Enrichment Analysis (see methods for details) are noted. LOD scores within the interval are included for the gene nearest to the marker position, with darkness of purple background indicating how close it is to the peak LOD score for the QTL; for those genes that fall between markers and do not have a specific LOD score, this color is estimated.

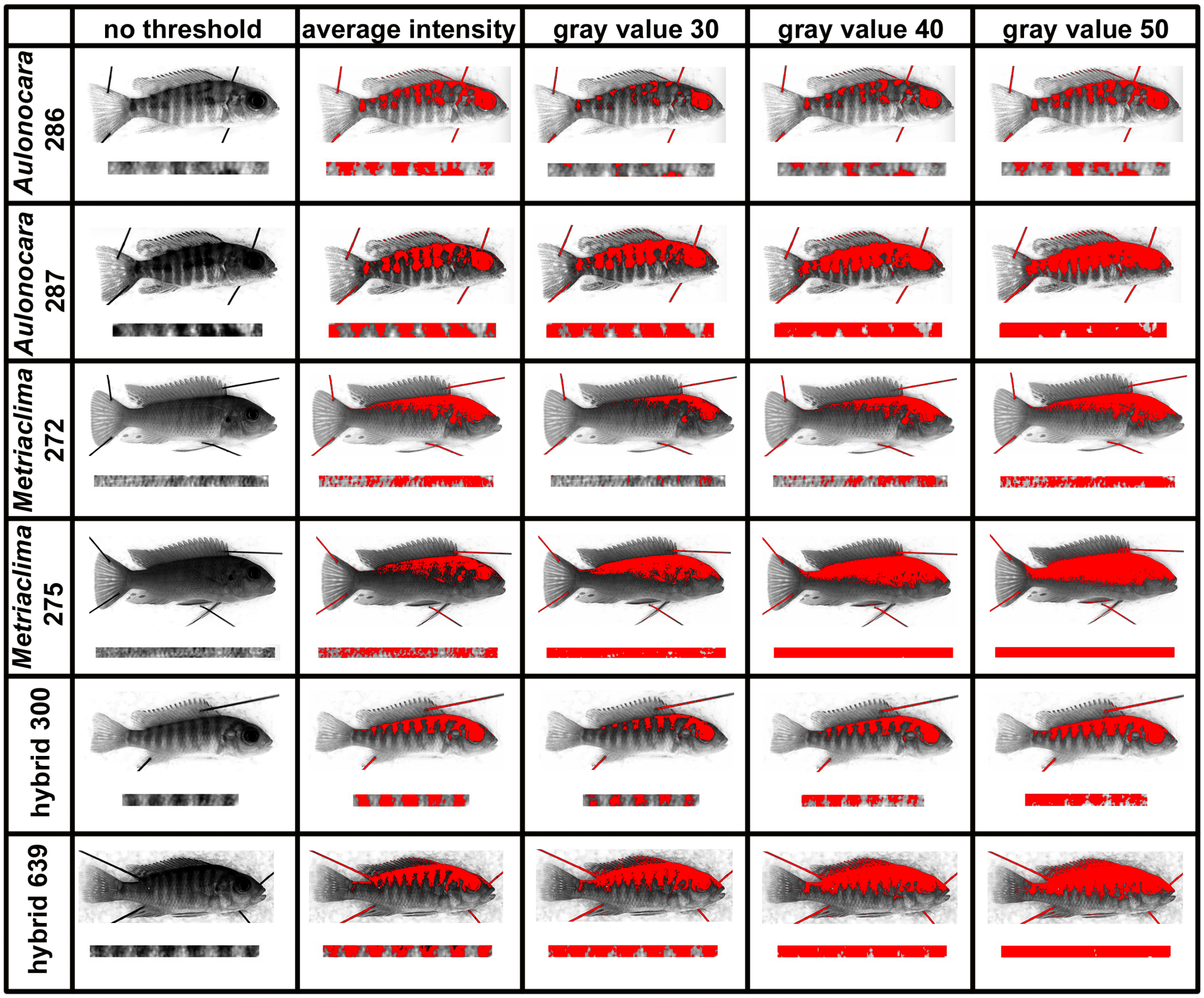


**Figure S1. Determination of thresholds used to define bars.** Empirical assessment was used to identify the appropriate gray intensity value to use as a cutoff to define bars and interbars. This was conducted on a test set of four *Aulonocara* parentals, four *Metriaclima* parentals, and four randomly-chosen hybrids, with two of each group visualized here. Each panel includes the picture of the entire animal on top and the isolated region used for analysis of barring. Pixels in red are those in which the pixel is equal to or less than (i.e., darker than) the indicated intensity, and are considered within a bar. Pixels with a gray intensity value greater (i.e., lighter) than the indicated cutoff is considered within an interbar. The average intensity measure represents the average value of the isolated region, calculated for each individual specimen. Note that using a cutoff of the average intensity for each individual most accurately represents the pattern of barring observed by eye.

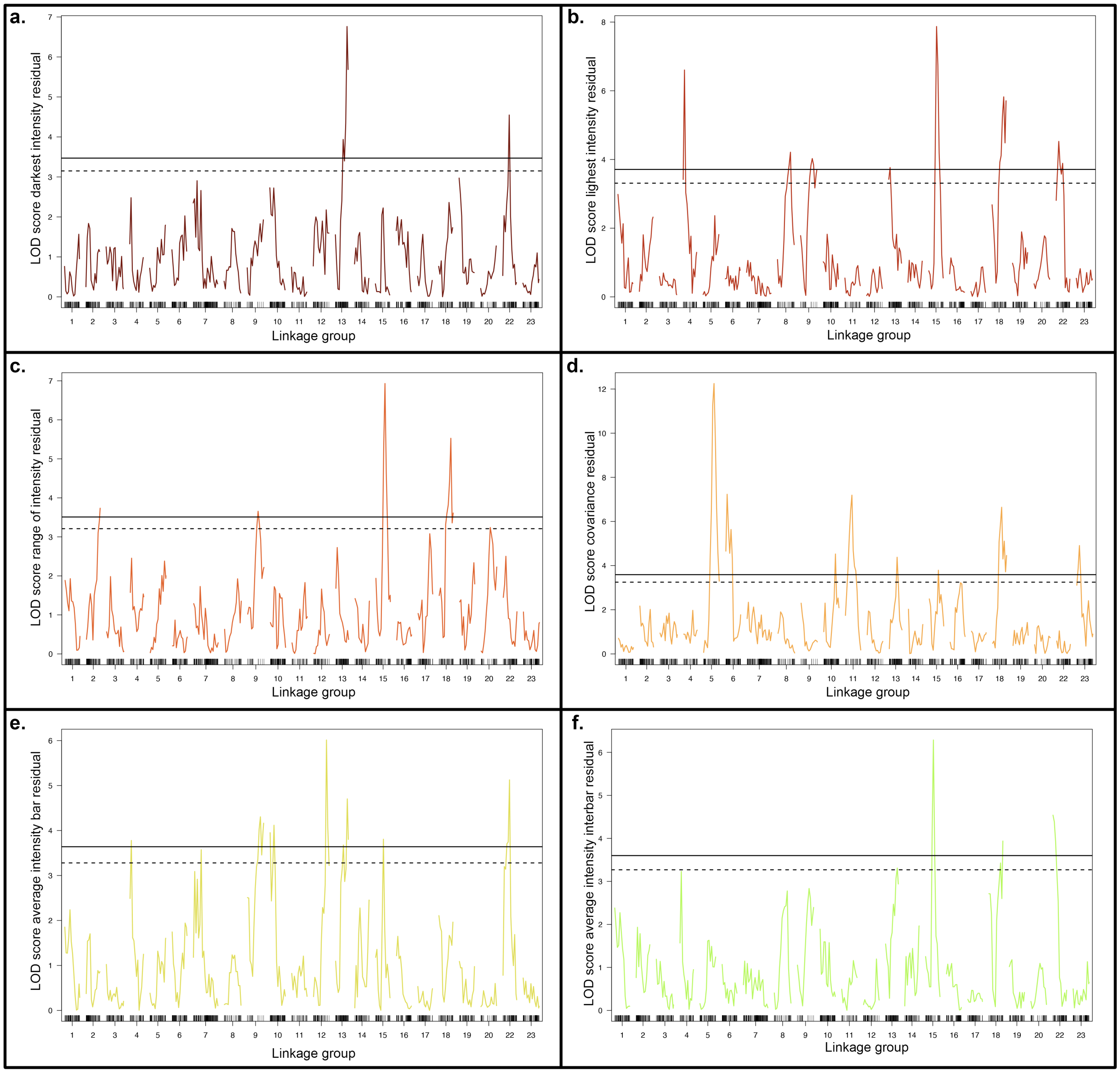
a.
LOD score darkest intensity residual
Linkage group
b.
LOD score lighest intensity residual
Linkage group
c.
LOD score range of intensity residual
Linkage group
d.
LOD score covariance residual
Linkage group
e.
LOD score average intensity bar residual
Linkage group
f.
LOD score average intensity interbar residual
Linkage group

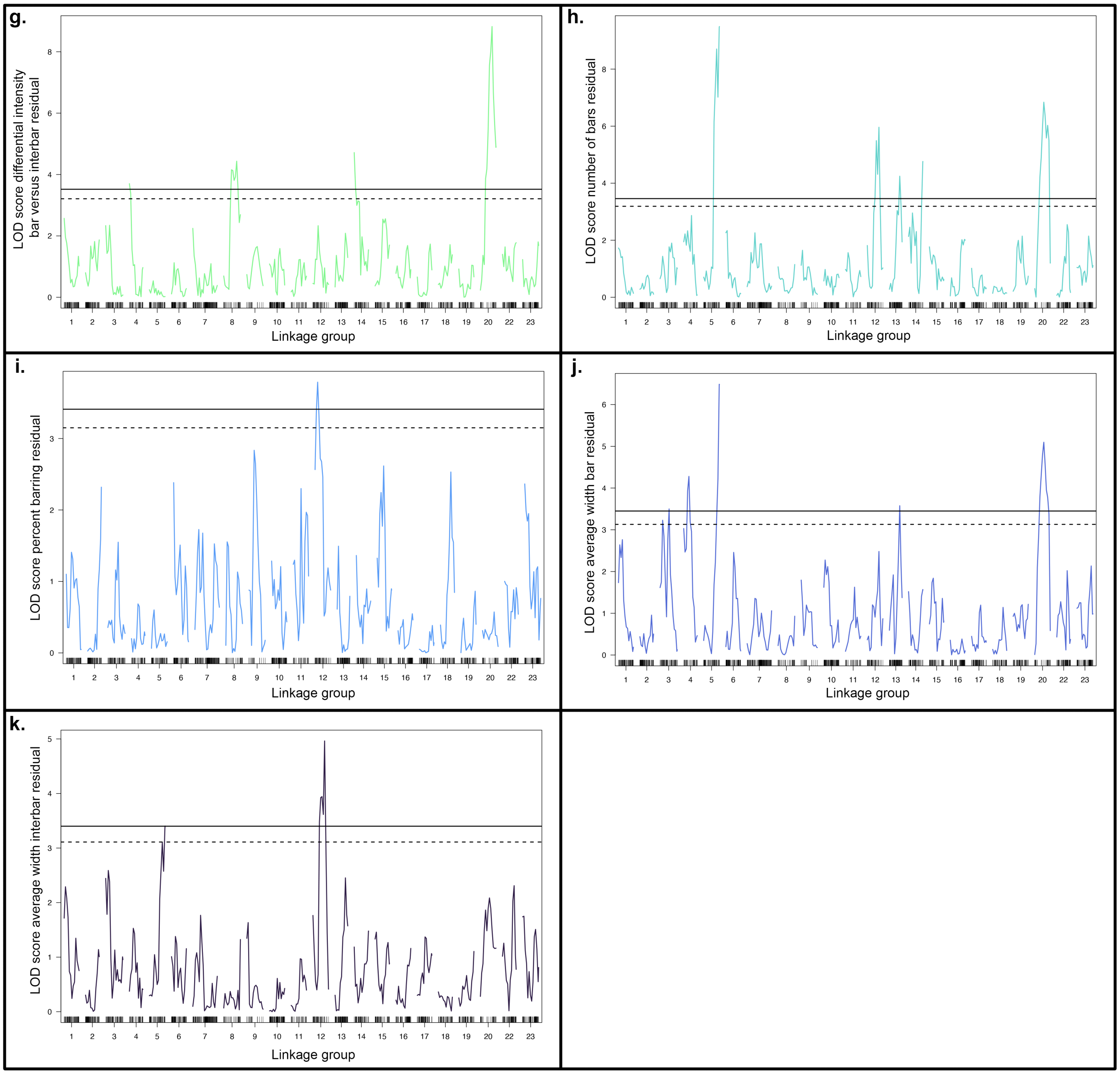


**Figure S2. Genome-wide QTL scans for pigment traits.** Pigment traits analyzed are residual data for (a) darkest intensity, (b) lightest intensity, (c) range of intensity, (d) covariance, (e) average intensity of bars, (f) average intensity of interbars, (g) differential intensity bars versus interbars, (h) number of bars, (i) percent barring, calculated as sum of total width of bars divided by total width of the isolated region, (j) average width of bars, and (k) average width of interbars. Colors of the scan match colors used in Figures 1 and 3. Significance is indicated at the 5% (solid line) and 10% (dashed line) level. Details of QTL scans are in Table S1. QTL scans by chromosome are in Figure S3.

a.

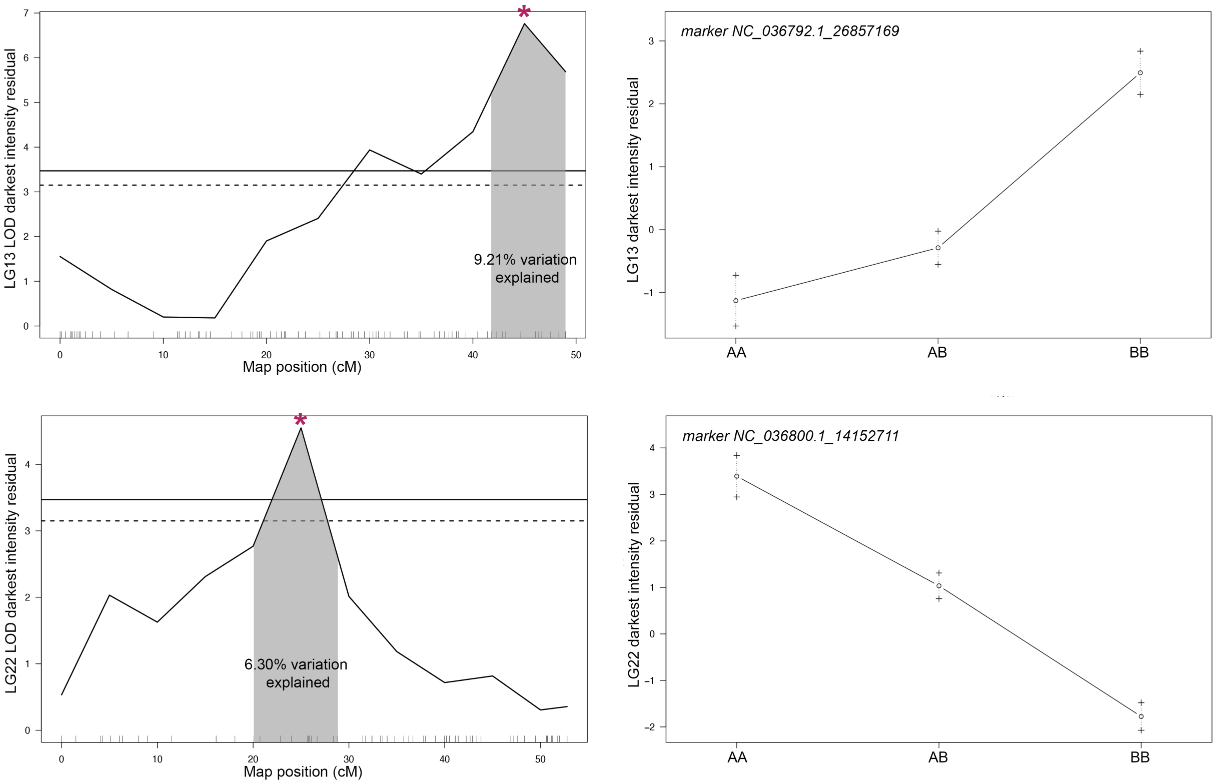
LG13 LOD darkest intensity residual
9.21% variation explained
Map position (cM)
marker NC_036792.1_26857169
LG13 darkest intensity residual
AA
AB
BB
LG22 LOD darkest intensity residual
6.30% variation explained
Map position (cM)
marker NC_036800.1_14152711
LG22 darkest intensity residual
AA
AB
BB

b.

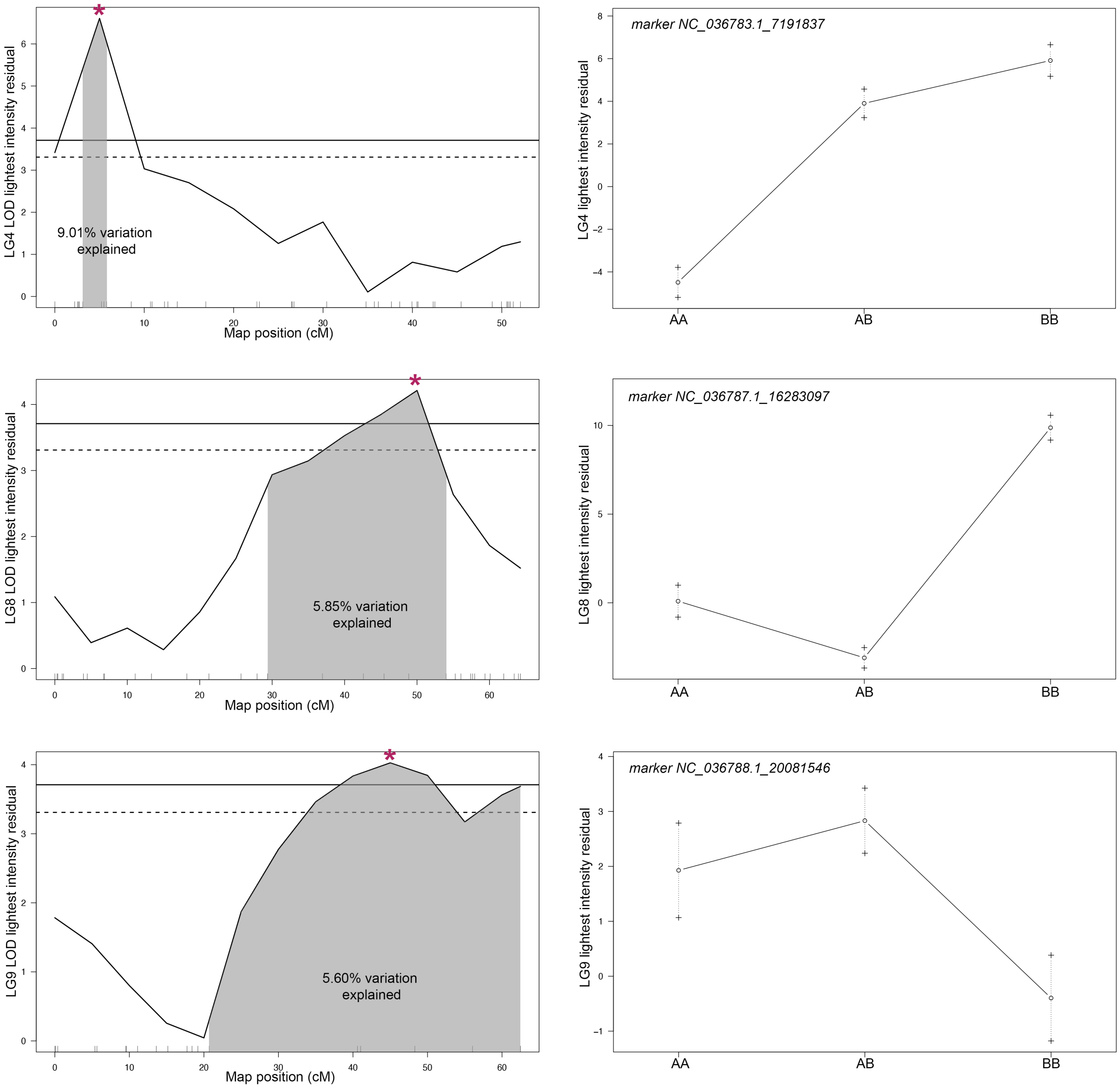

9.01% variation explained
LG4 LOD lightest intensity residual
Map position (cM)
marker NC_036783.1_7191837
LG4 lightest intensity residual
AA
AB
BB
5.85% variation explained
LG8 LOD lightest intensity residual
Map position (cM)
marker NC_036787.1_16283097
LG8 lightest intensity residual
AA
AB
BB
5.60% variation explained
LG9 LOD lightest intensity residual
Map position (cM)
marker NC_036788.1_20081546
LG9 lightest intensity residual
AA
AB
BB

**b.**

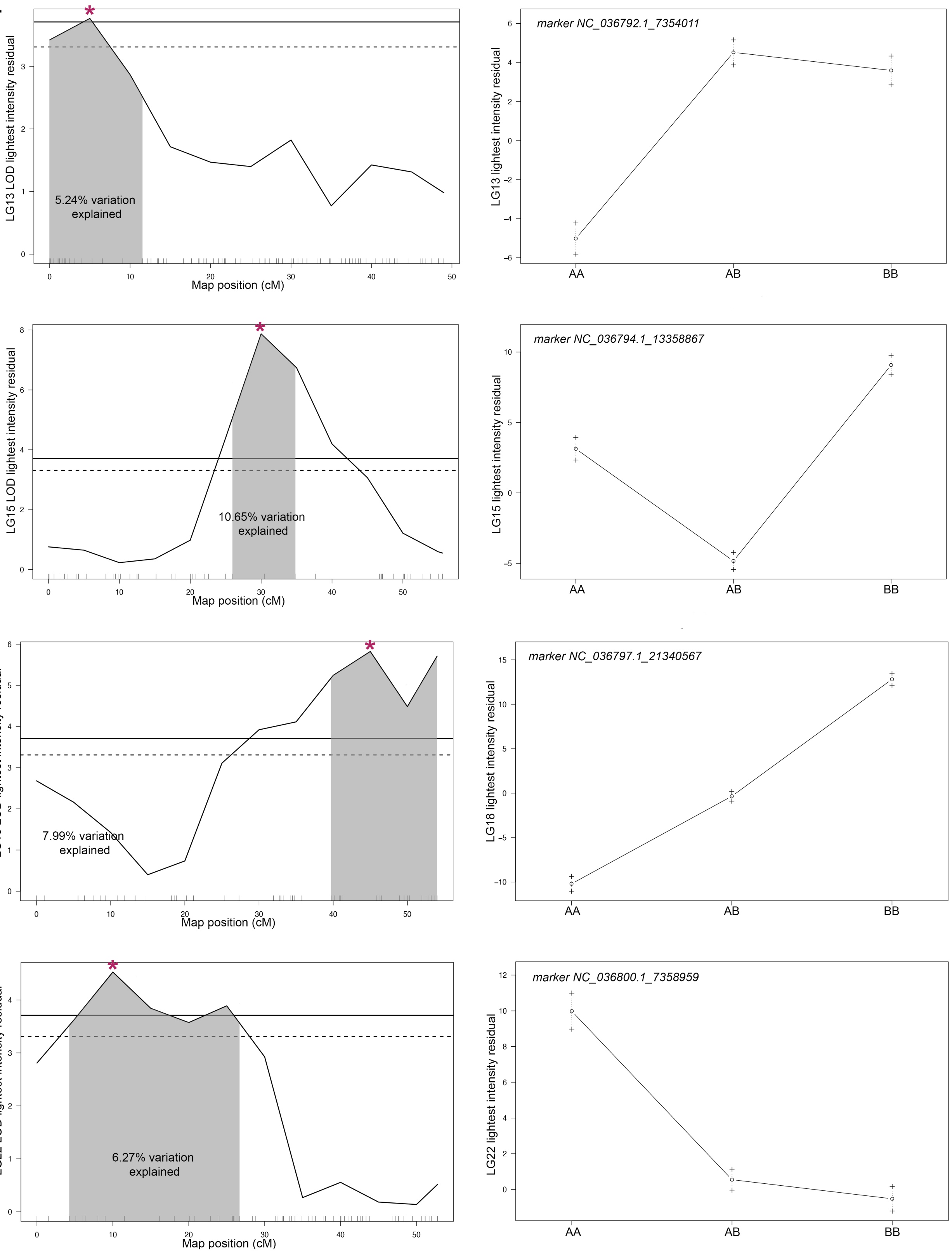

LG13 LOD lightest intensity residual
5.24% variation explained
Map position (cM)
marker NC_036792.1_7354011
LG13 lightest intensity residual
AA
AB
BB
LG15 LOD lightest intensity residual
10.65% variation explained
Map position (cM)
marker NC_036794.1_13358867
LG15 lightest intensity residual
AA
AB
BB
LG18 LOD lightest intensity residual
7.99% variation explained
Map position (cM)
marker NC_036797.1_21340567
LG18 lightest intensity residual
AA
AB
BB
LG22 LOD lightest intensity residual
6.27% variation explained
Map position (cM)
marker NC_036800.1_7358959
LG22 lightest intensity residual
AA
AB
BB

c.

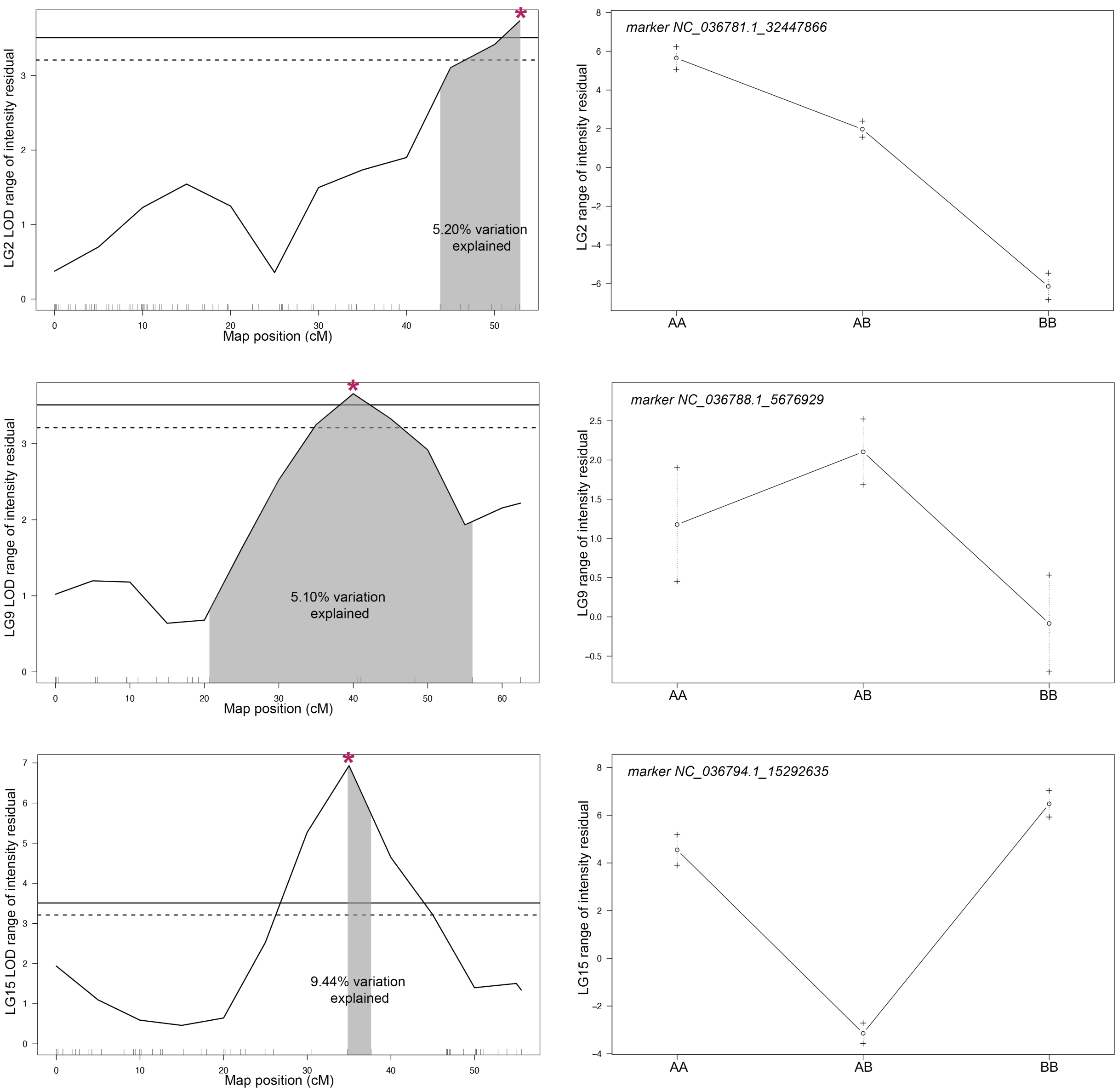
LG2 LOD range of intensity residual
5.20% variation explained
Map position (cM)
marker NC_036781.1_32447866
LG2 range of intensity residual
AA
AB
BB
LG9 LOD range of intensity residual
5.10% variation explained
Map position (cM)
marker NC_036788.1_5676929
LG9 range of intensity residual
AA
AB
BB
LG15 LOD range of intensity residual
9.44% variation explained
Map position (cM)
marker NC_036794.1_15292635
LG15 range of intensity residual
AA
AB
BB

c.

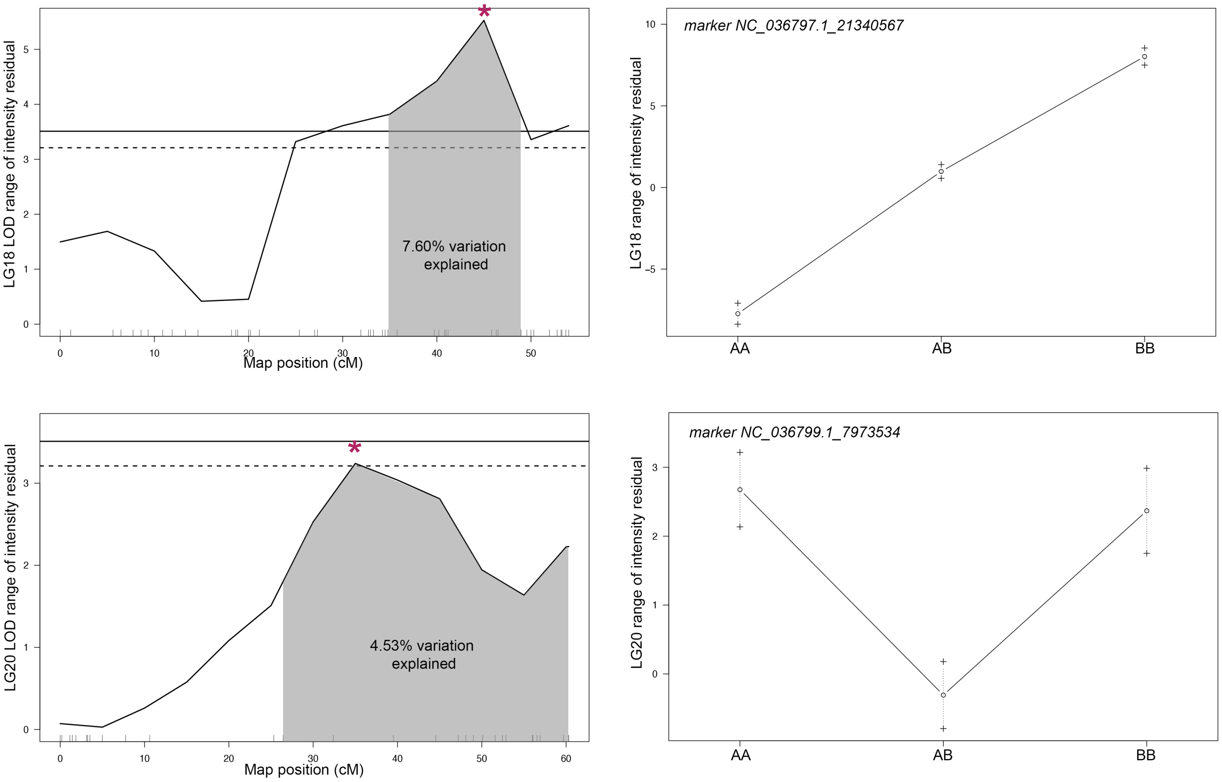

LG18 LOD range of intensity residual
7.60% variation explained
Map position (cM)
marker NC_036797.1_21340567
LG18 range of intensity residual
AA
AB
BB
LG20 LOD range of intensity residual
4.53% variation explained
Map position (cM)
marker NC_036799.1_7973534
LG20 range of intensity residual
AA
AB
BB

d.

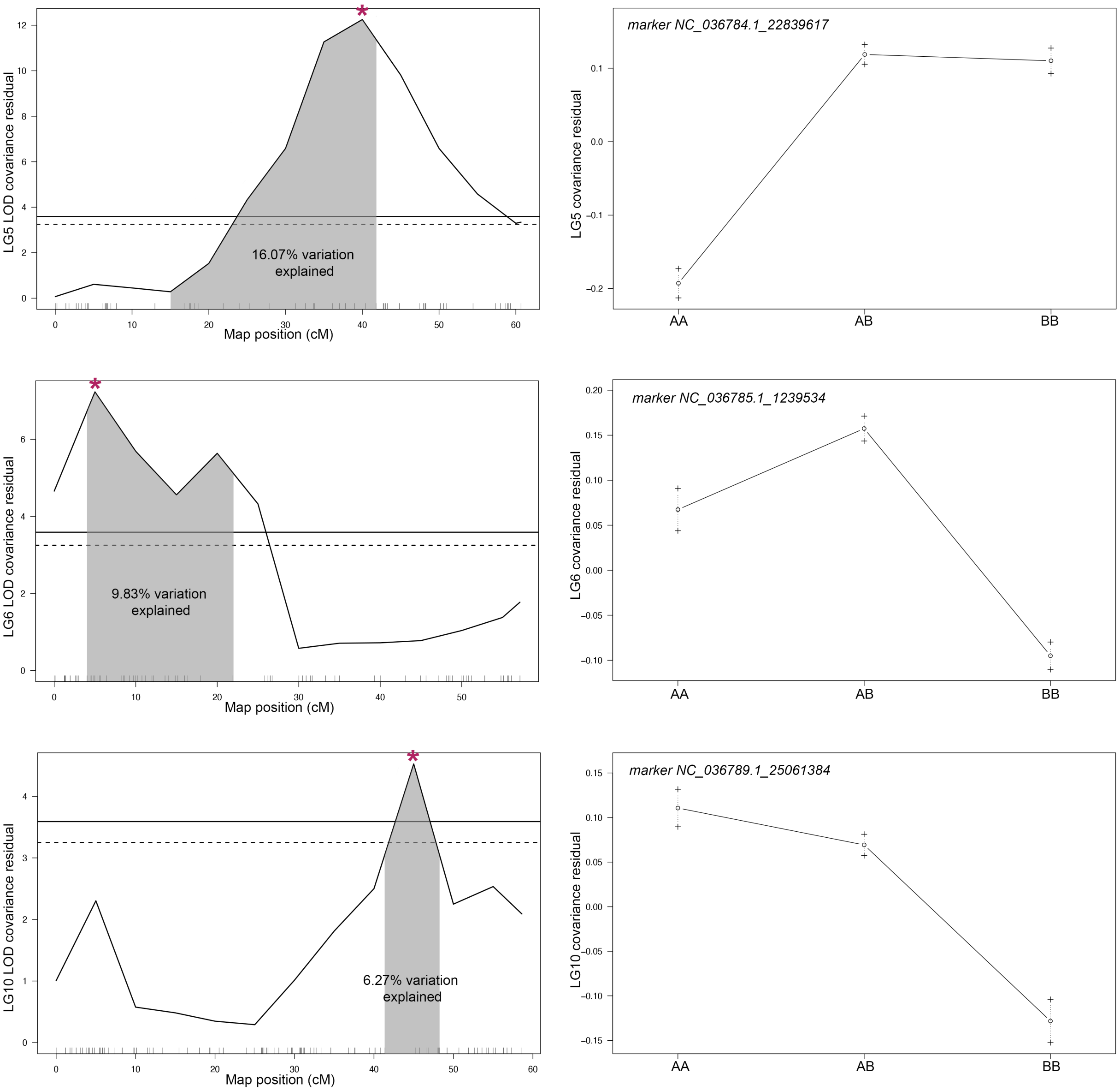
LG5 LOD covariance residual
16.07% variation
explained
Map position (cM)
marker NC_036784.1_22839617
LG5 covariance residual
AA
AB
BB
LG6 LOD covariance residual
9.83% variation
explained
Map position (cM)
marker NC_036785.1_1239534
LG6 covariance residual
AA
AB
BB
LG10 LOD covariance residual
6.27% variation
explained
Map position (cM)
marker NC_036789.1_25061384
LG10 covariance residual
AA
AB
BB

d.

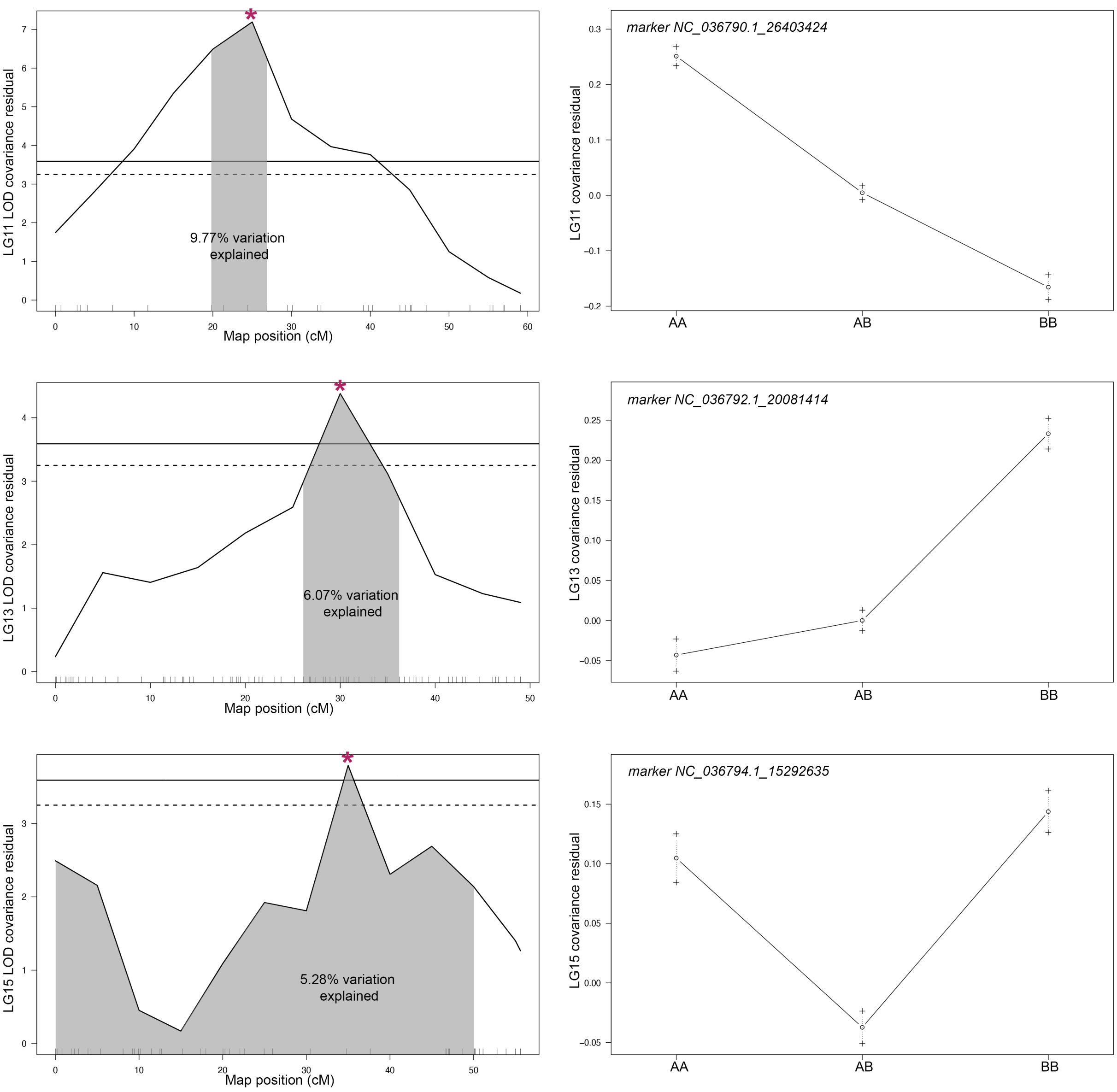
LG11 LOD covariance residual
9.77% variation explained
Map position (cM)
marker NC_036790.1_26403424
LG11 covariance residual
AA
AB
BB
LG13 LOD covariance residual
6.07% variation explained
Map position (cM)
marker NC_036792.1_20081414
LG13 covariance residual
AA
AB
BB
LG15 LOD covariance residual
5.28% variation explained
Map position (cM)
marker NC_036794.1_15292635
LG15 covariance residual
AA
AB
BB

d.

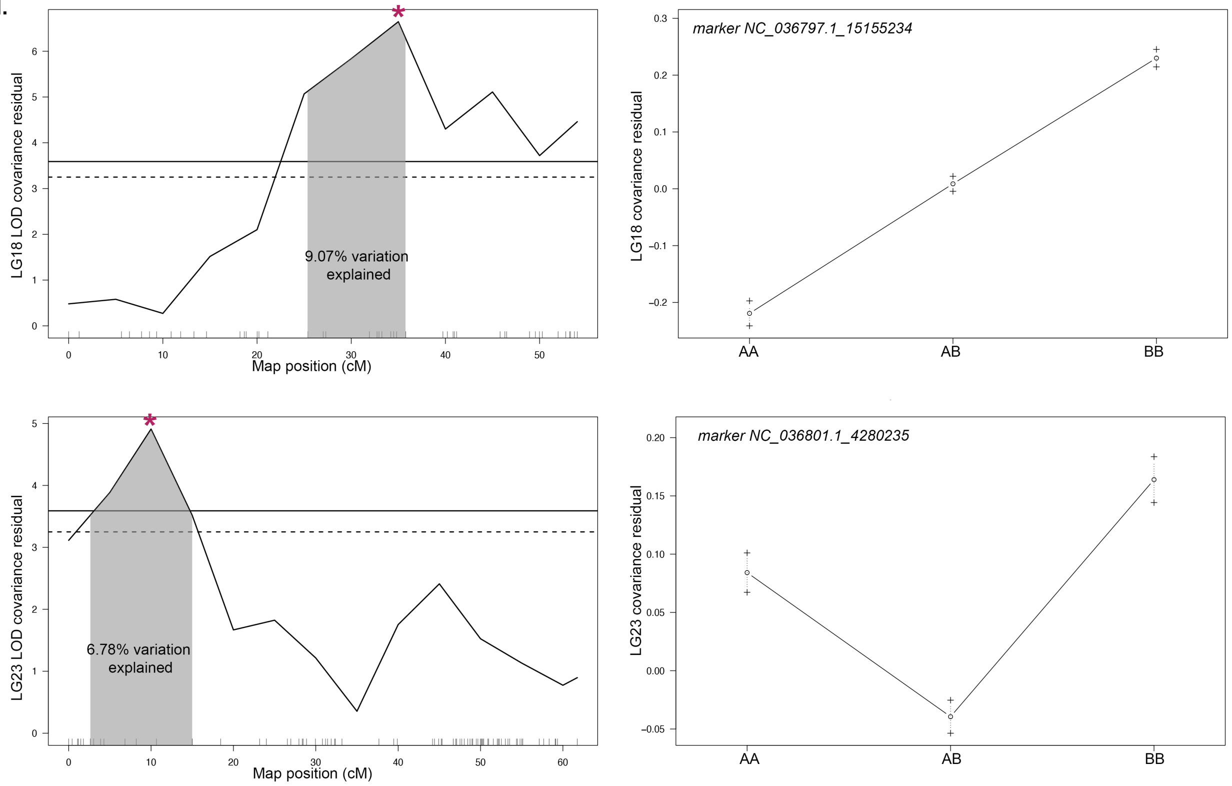
LG18 LOD covariance residual
9.07% variation explained
Map position (cM)
marker NC_036797.1_15155234
LG18 covariance residual
AA
AB
BB
LG23 LOD covariance residual
6.78% variation explained
Map position (cM)
marker NC_036801.1_4280235
LG23 covariance residual
AA
AB
BB

e.

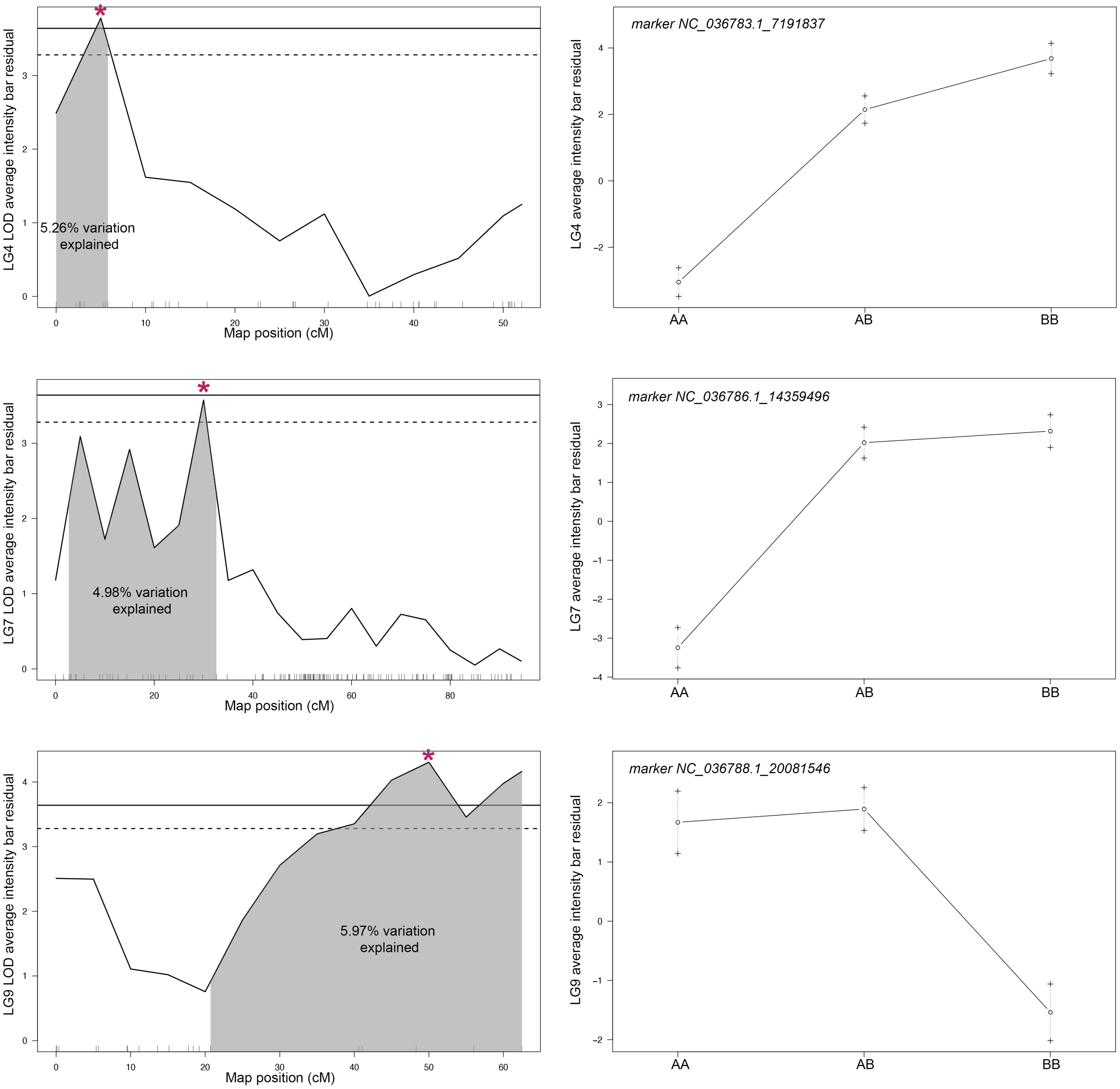

5.26% variation explained
LG4 LOD average intensity bar residual
Map position (cM)
marker NC_036783.1_7191837
LG4 average intensity bar residual
AA
AB
BB
4.98% variation explained
LG7 LOD average intensity bar residual
Map position (cM)
marker NC_036786.1_14359496
LG7 average intensity bar residual
AA
AB
BB
5.97% variation explained
LG9 LOD average intensity bar residual
Map position (cM)
marker NC_036788.1_20081546
LG9 average intensity bar residual
AA
AB
BB

**e.**

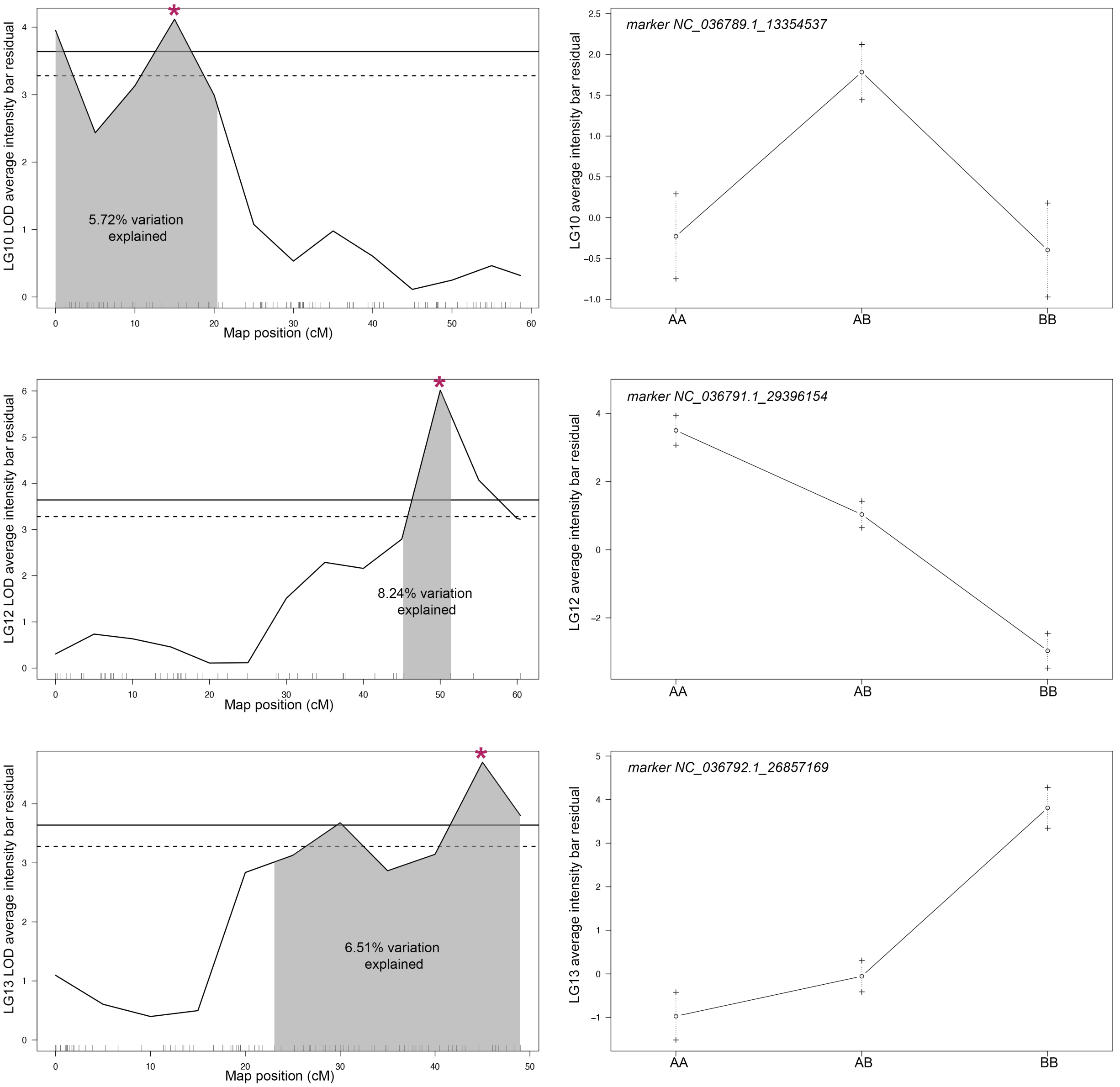
LG10 LOD average intensity bar residual
5.72% variation
explained
Map position (cM)
marker NC_036789.1_13354537
LG10 average intensity bar residual
AA
AB
BB
LG12 LOD average intensity bar residual
8.24% variation
explained
Map position (cM)
marker NC_036791.1_29396154
LG12 average intensity bar residual
AA
AB
BB
LG13 LOD average intensity bar residual
6.51% variation
explained
Map position (cM)
marker NC_036792.1_26857169
LG13 average intensity bar residual
AA
AB
BB

**e.**

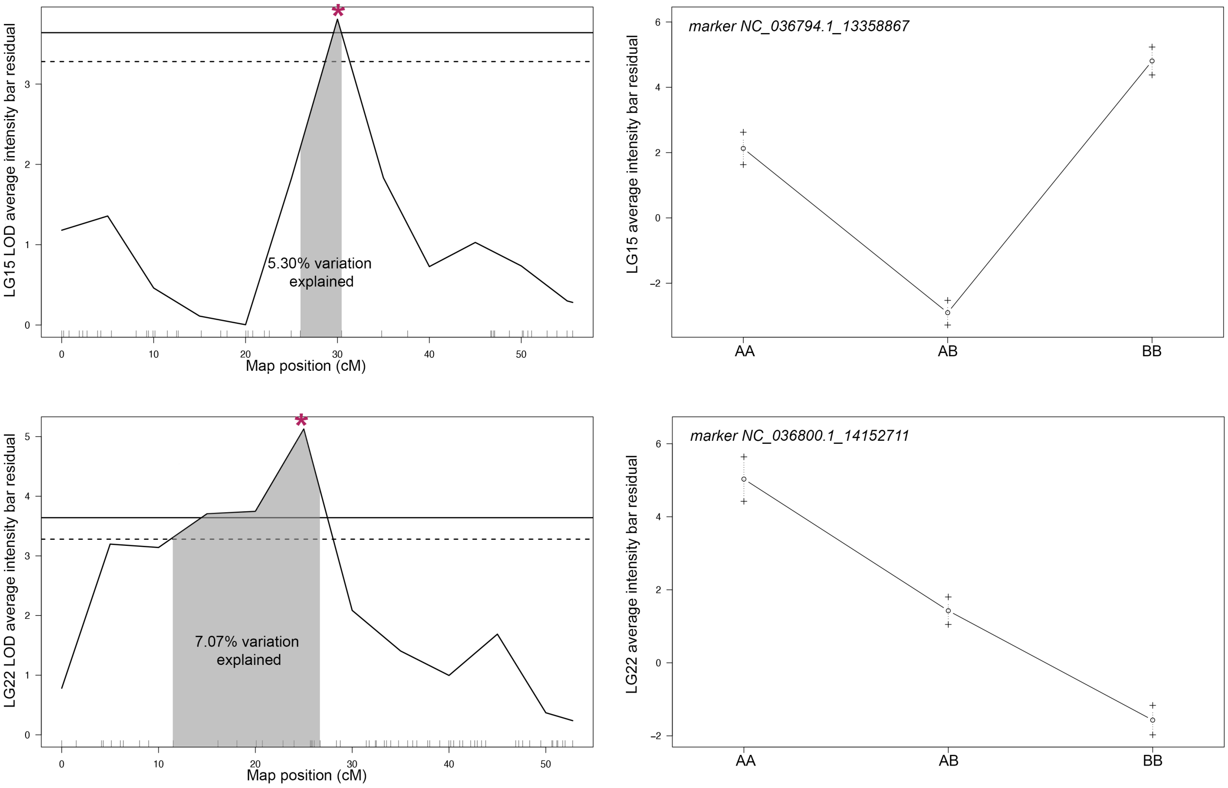

LG15 LOD average intensity bar residual
5.30% variation explained
Map position (cM)
marker NC_036794.1_13358867
LG15 average intensity bar residual
AA
AB
BB
LG22 LOD average intensity bar residual
7.07% variation explained
Map position (cM)
marker NC_036800.1_14152711
LG22 average intensity bar residual
AA
AB
BB


**f.**

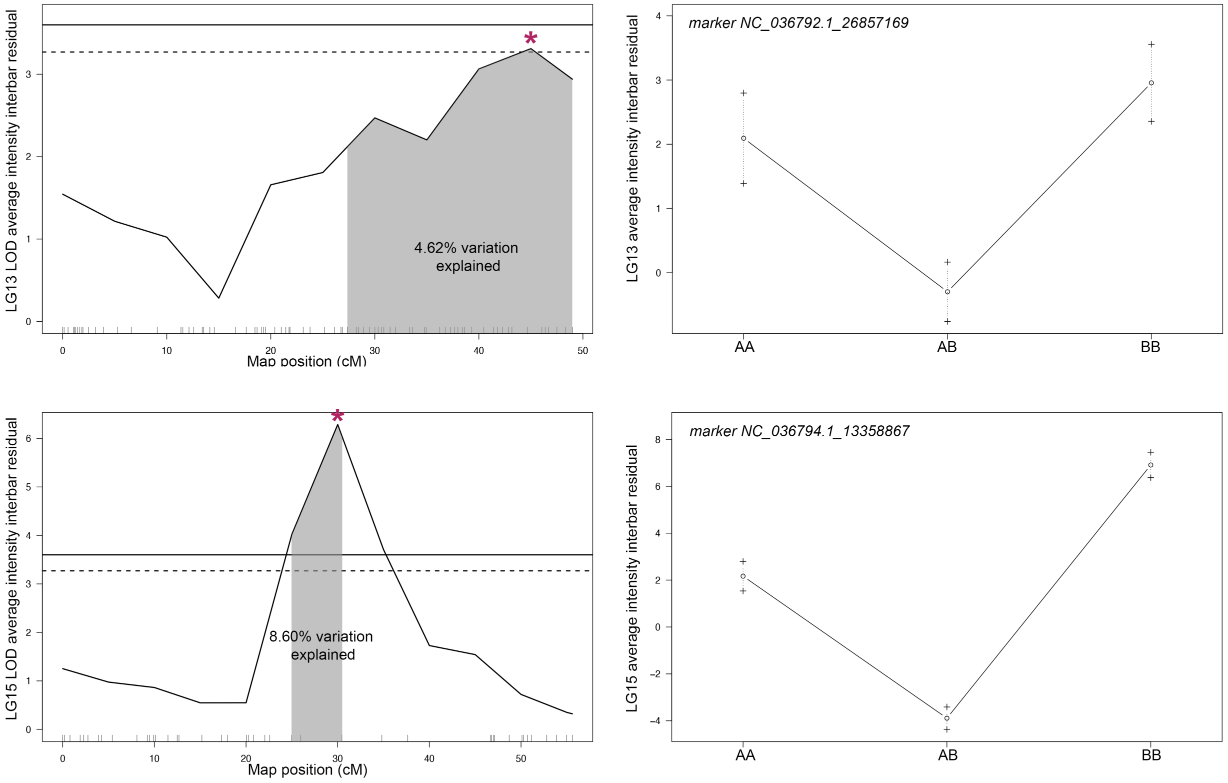

LG13 LOD average intensity interbar residual
4.62% variation explained
Map position (cM)
marker NC_036792.1_26857169
LG13 average intensity interbar residual
AA
AB
BB
LG15 LOD average intensity interbar residual
8.60% variation explained
Map position (cM)
marker NC_036794.1_13358867
LG15 average intensity interbar residual
AA
AB
BB

f.

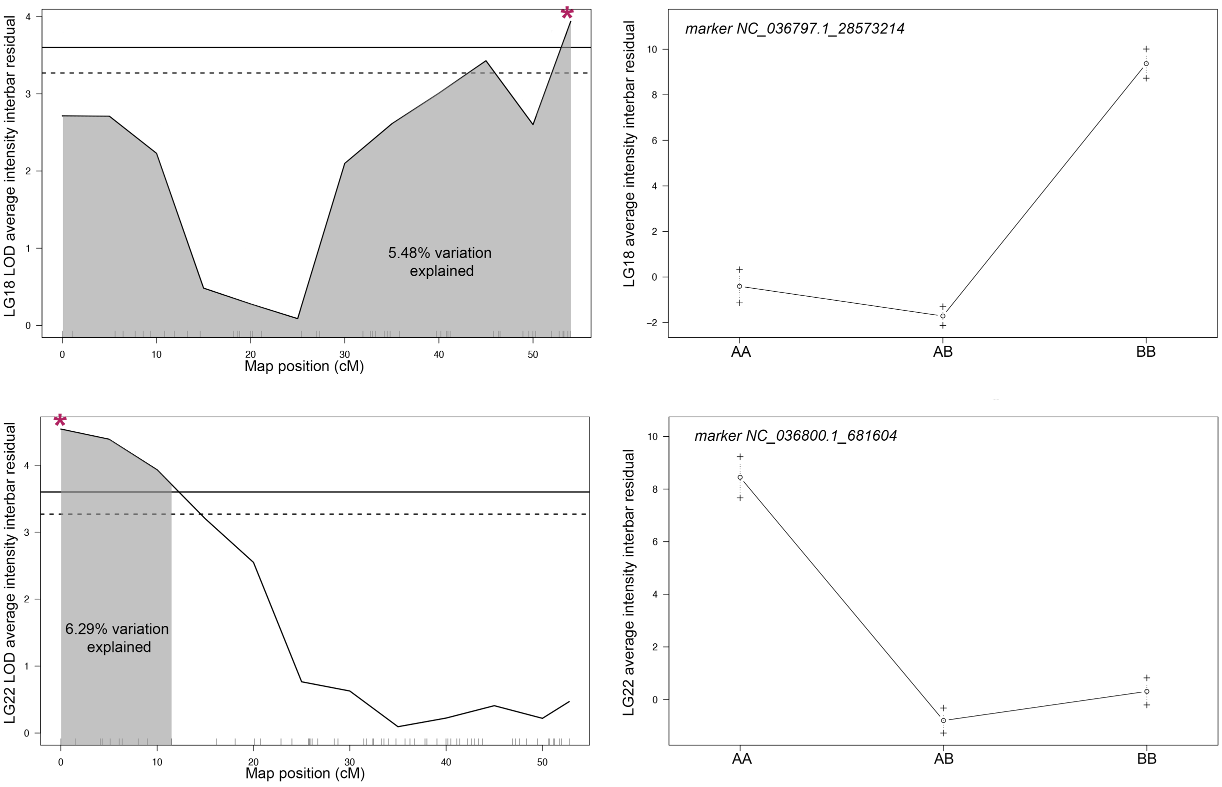
LG18 LOD average intensity interbar residual
5.48% variation explained
Map position (cM)
marker NC_036797.1_28573214
LG18 average intensity interbar residual
AA
AB
BB
LG22 LOD average intensity interbar residual
6.29% variation explained
Map position (cM)
marker NC_036800.1_681604
LG22 average intensity interbar residual
AA
AB
BB

g.

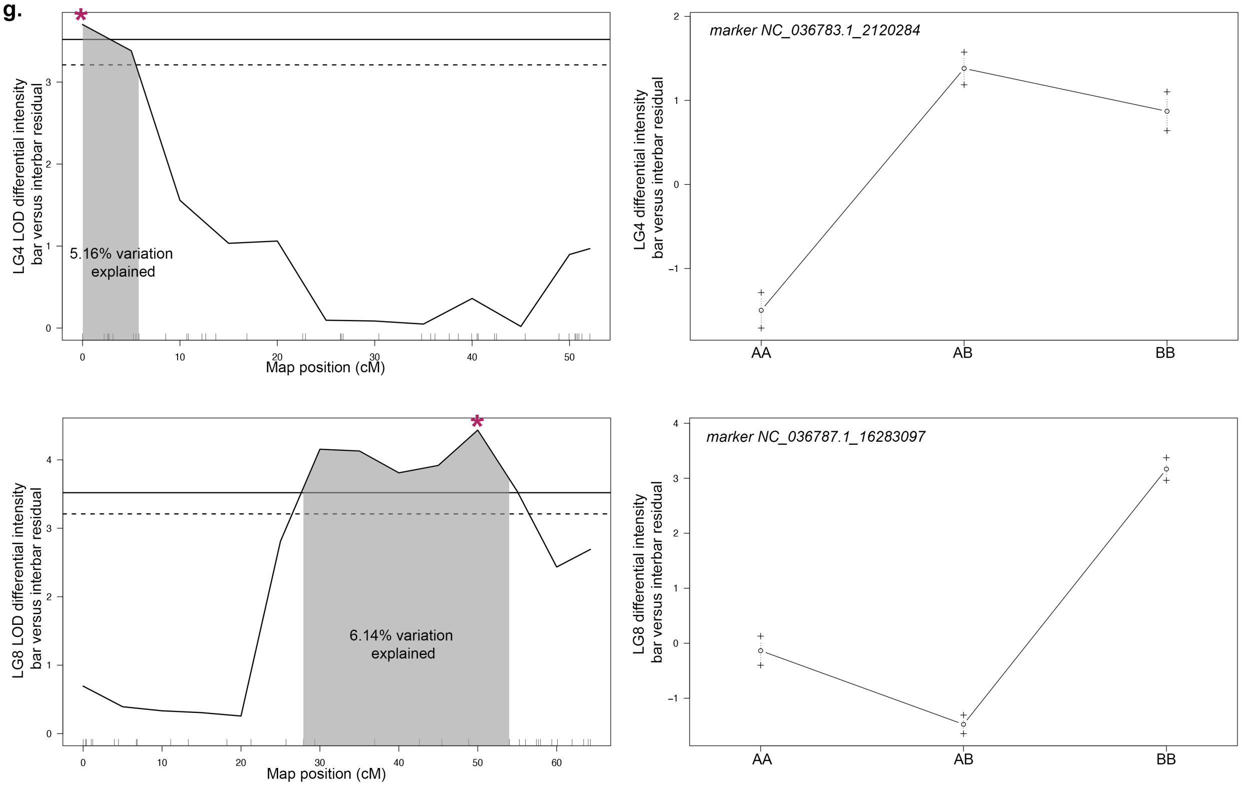
LG4 LOD differential intensity bar versus interbar residual
5.16% variation explained
Map position (cM)
marker NC_036783.1_2120284
LG4 differential intensity bar versus interbar residual
AA
AB
BB
LG8 LOD differential intensity bar versus interbar residual
6.14% variation explained
Map position (cM)
marker NC_036787.1_16283097
LG8 differential intensity bar versus interbar residual
AA
AB
BB

g.

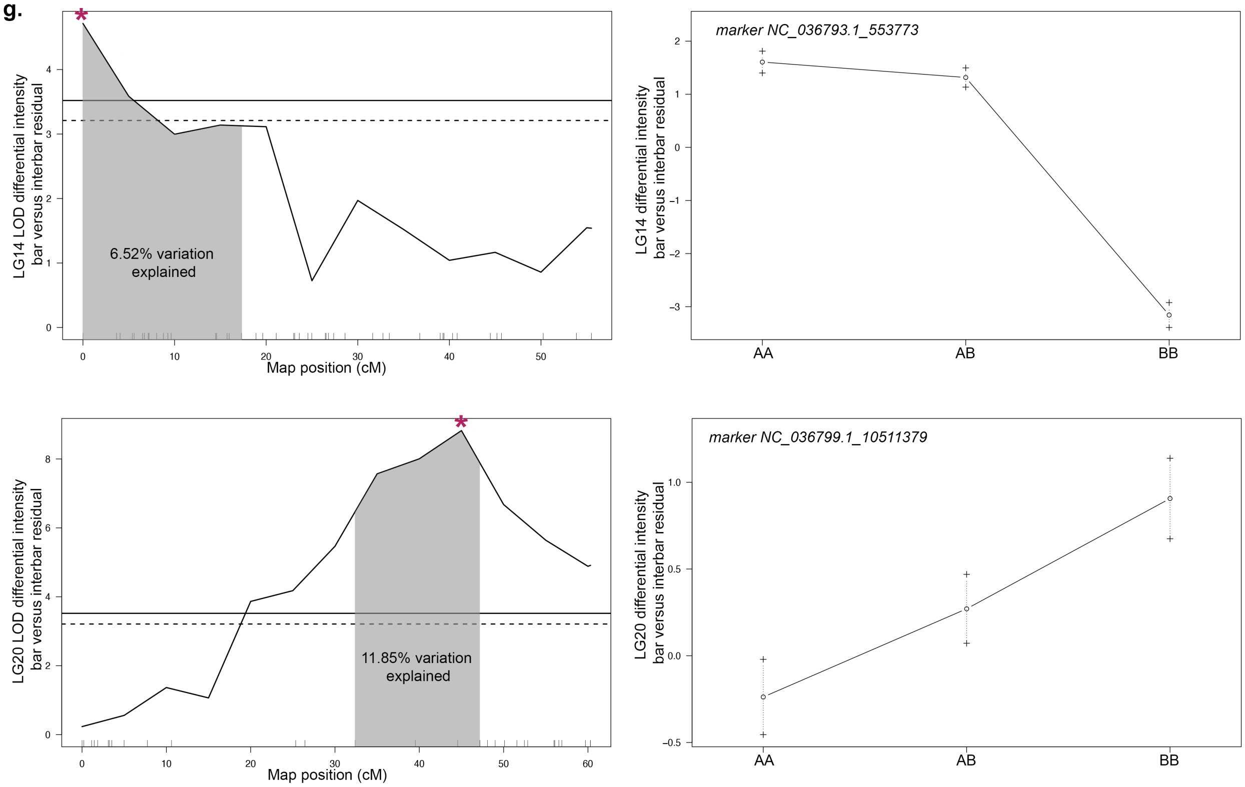

LG14 LOD differential intensity bar versus interbar residual
6.52% variation explained
Map position (cM)
marker NC_036793.1_553773
LG14 differential intensity bar versus interbar residual
AA
AB
BB
LG20 LOD differential intensity bar versus interbar residual
11.85% variation explained
Map position (cM)
marker NC_036799.1_10511379
LG20 differential intensity bar versus interbar residual
AA
AB
BB

**h.**

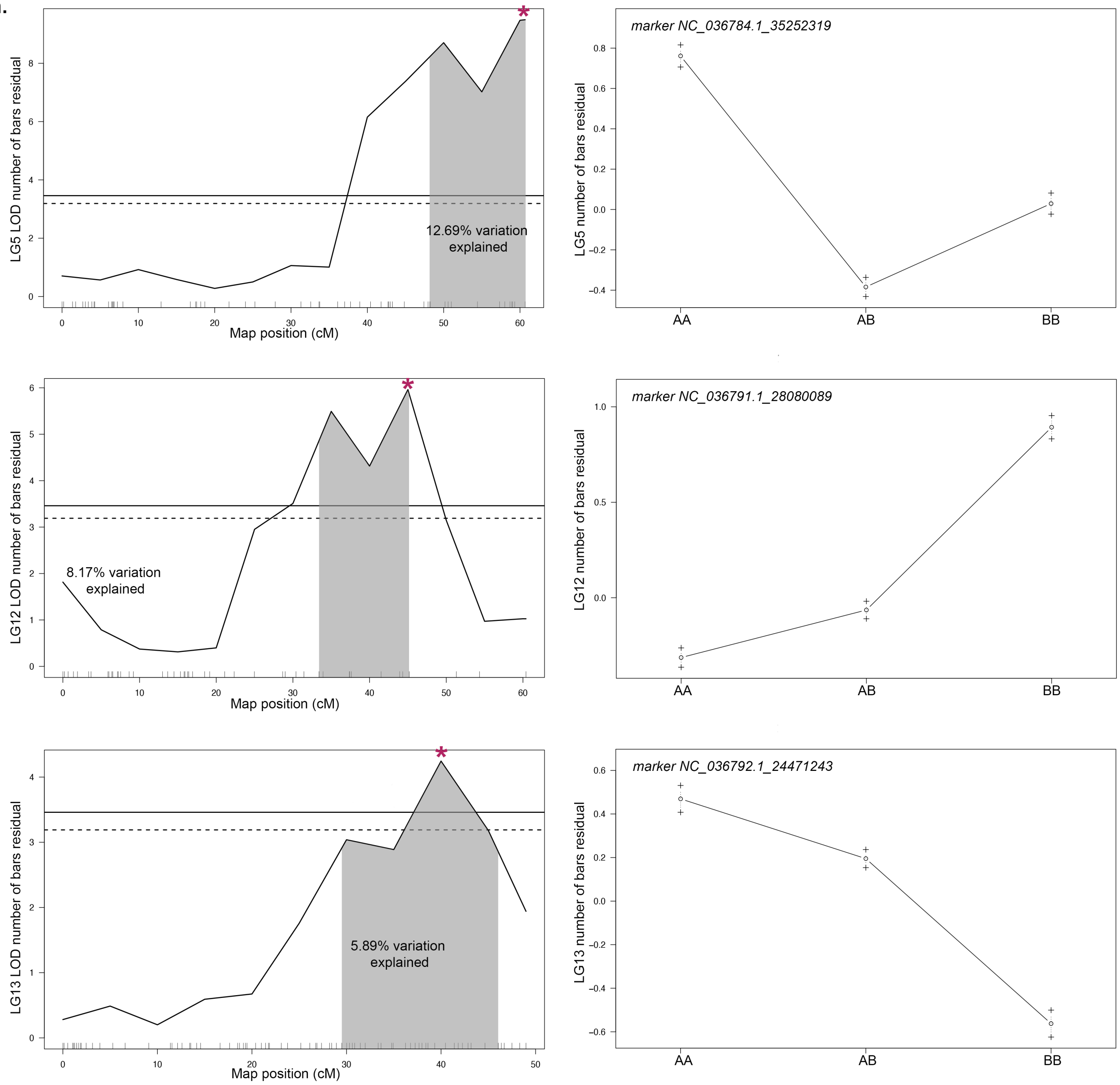
LG5 LOD number of bars residual
Map position (cM)
12.69% variation explained
marker NC_036784.1_35252319
LG5 number of bars residual
AA
AB
BB
LG12 LOD number of bars residual
Map position (cM)
8.17% variation explained
marker NC_036791.1_28080089
LG12 number of bars residual
AA
AB
BB
LG13 LOD number of bars residual
Map position (cM)
5.89% variation explained
marker NC_036792.1_24471243
LG13 number of bars residual
AA
AB
BB

h.

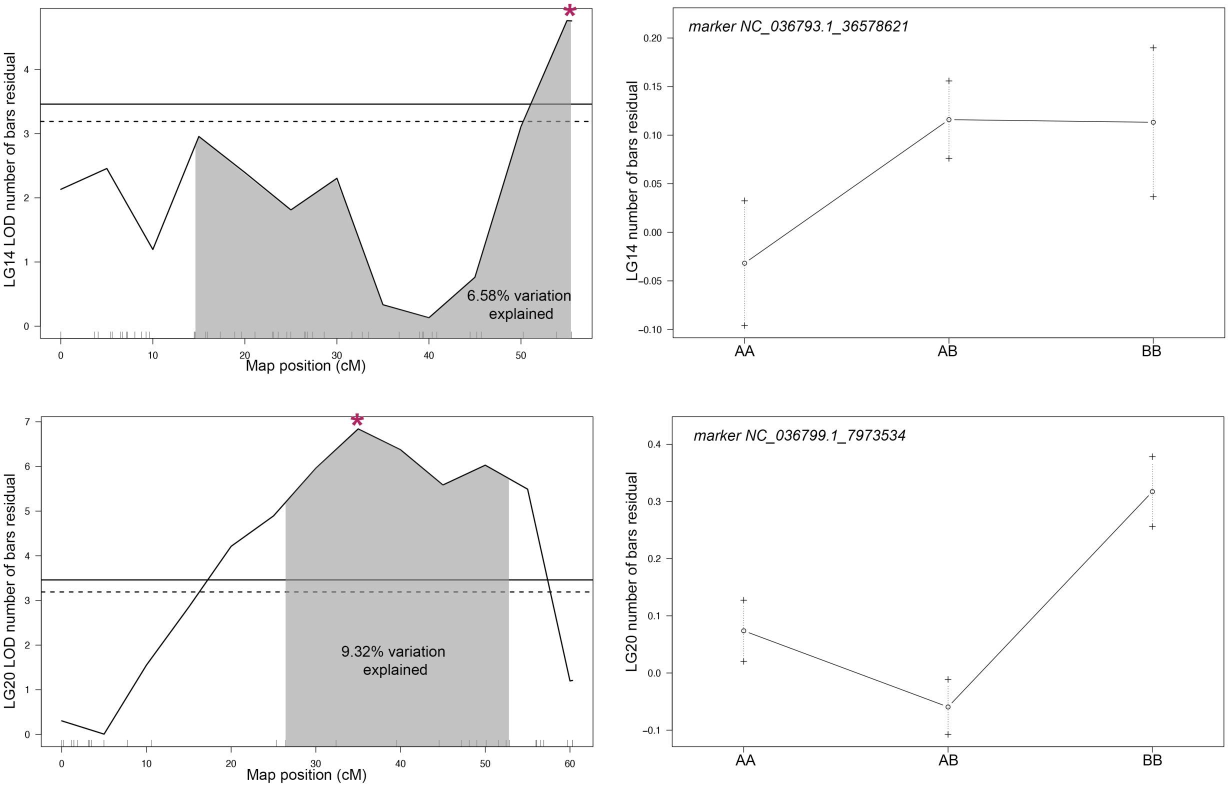
LG14 LOD number of bars residual
6.58% variation explained
Map position (cM)
marker NC_036793.1_36578621
LG14 number of bars residual
AA
AB
BB
LG20 LOD number of bars residual
9.32% variation explained
Map position (cM)
marker NC_036799.1_7973534
LG20 number of bars residual
AA
AB
BB

i.

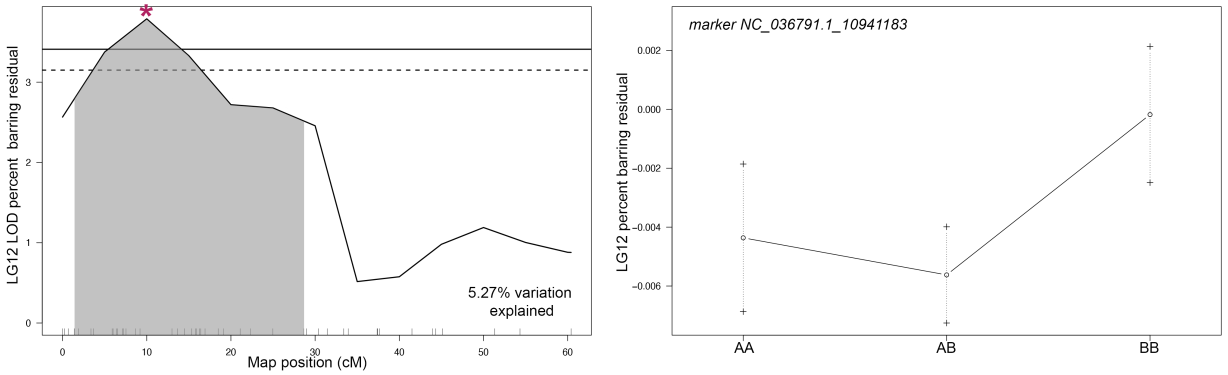
LG12 LOD percent barring residual
5.27% variation explained
Map position (cM)
marker NC_036791.1_10941183
LG12 percent barring residual
AA
AB
BB

**j.**

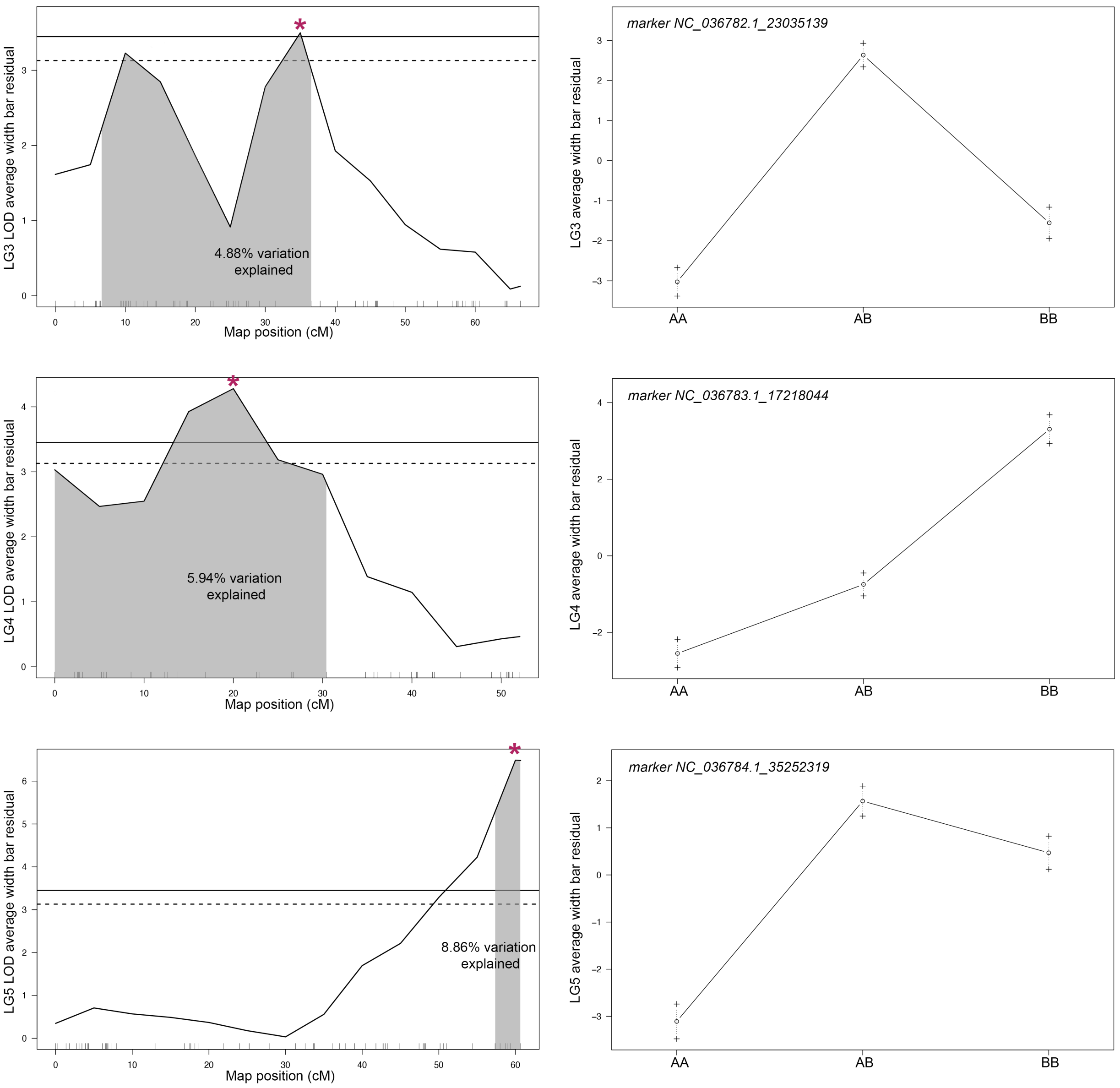
*
LG3 LOD average width bar residual
4.88% variation explained
Map position (cM)
marker NC_036782.1_23035139
LG3 average width bar residual
AA
AB
BB
*
LG4 LOD average width bar residual
5.94% variation explained
Map position (cM)
marker NC_036783.1_17218044
LG4 average width bar residual
AA
AB
BB
*
LG5 LOD average width bar residual
8.86% variation explained
Map position (cM)
marker NC_036784.1_35252319
LG5 average width bar residual
AA
AB
BB

j.

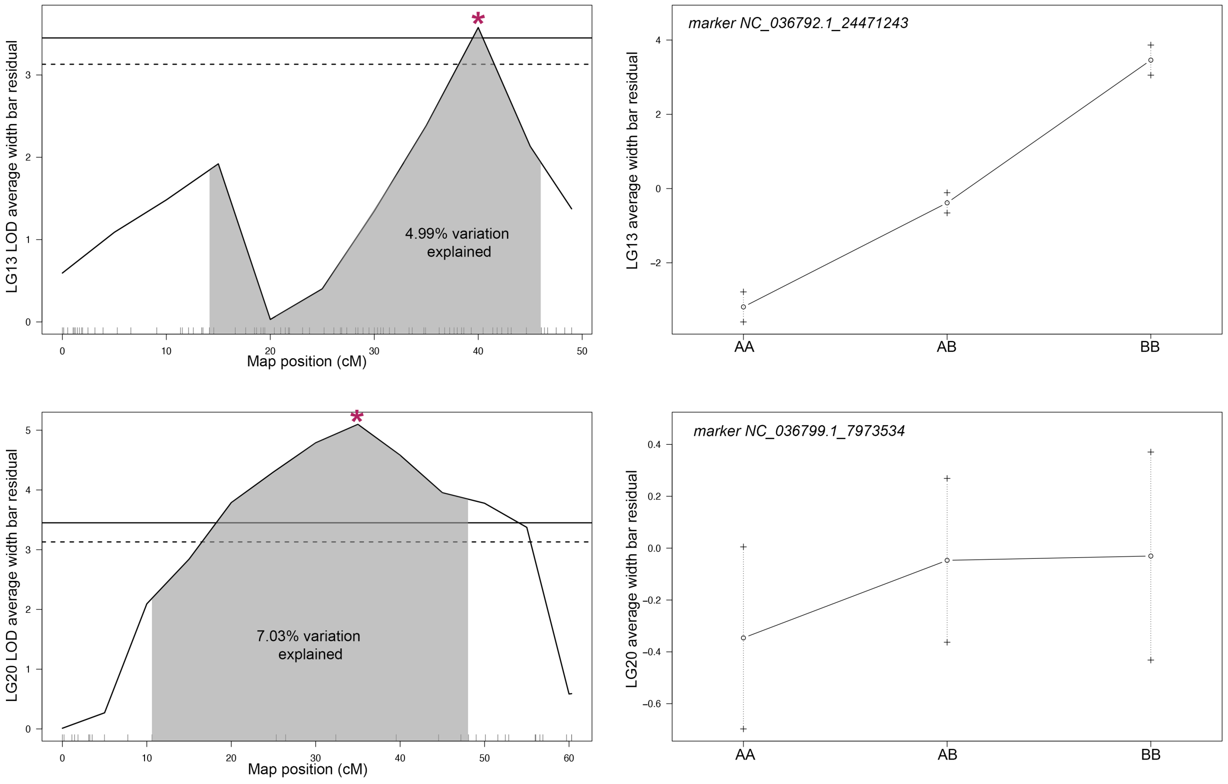
LG13 LOD average width bar residual
Map position (cM)
4.99% variation explained
marker NC_036792.1_24471243
LG13 average width bar residual
AA
AB
BB
LG20 LOD average width bar residual
Map position (cM)
7.03% variation explained
marker NC_036799.1_7973534
LG20 average width bar residual
AA
AB
BB

k.

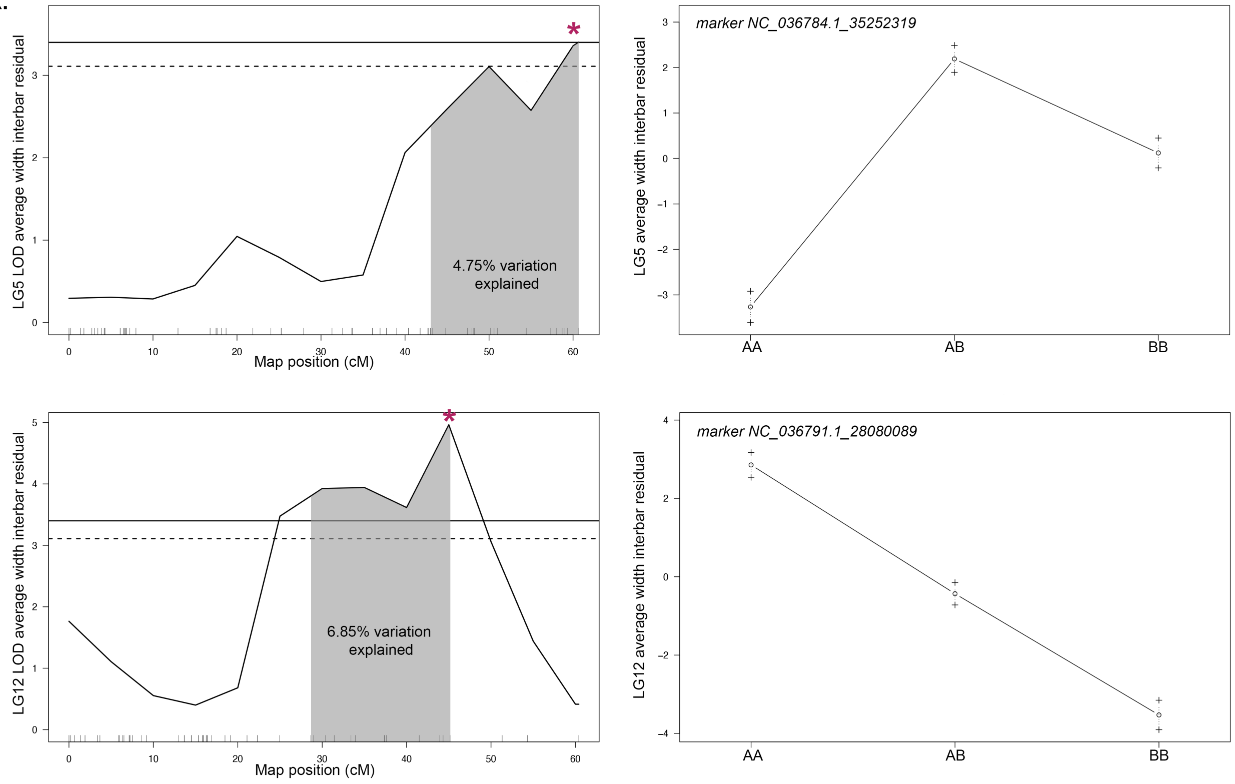
LG5 LOD average width interbar residual
Map position (cM)
4.75% variation explained
marker NC_036784.1_35252319
LG5 average width interbar residual
AA
AB
BB
LG12 LOD average width interbar residual
Map position (cM)
6.85% variation explained
marker NC_036791.1_28080089
LG12 average width interbar residual
AA
AB
BB

**Figure S3. QTL scans by linkage group and allelic effects.** Pigment traits analyzed are residual data for (a) darkest intensity, (b) lightest intensity, (c) range of intensity, (d) covariance, (e) average intensity of bars, (f) average intensity of interbars, (g) differential intensity bars versus interbars, (h) number of bars, (i) percent barring, calculated as sum of total width of bars divided by total width of the isolated region, (j) average width of bars, and (k) average width of interbars. 95% confidence interval for QTL is indicated by shading, percent of total phenotypic variation explained by QTL is reported, and genome-wide significance is shown at the 5% (solid line) and 10% (dashed line) level. Details of QTL scan are in Table S1 and genome-wide visuals are in Figure S2. Allelic effects are shown for marker at the peak log odds (LOD) score for the QTL, which is indicated by *. The A allele was inherited from the *Metriaclima* granddam and the B allele from the *Aulonocara* grandsire.
